# *morphoHeart*: a novel quantitative tool to perform integrated 3D morphometric analyses of heart and ECM morphology during embryonic development

**DOI:** 10.1101/2024.02.19.580991

**Authors:** Juliana Sánchez-Posada, Emily S Noël

**Affiliations:** School of Biosciences and Bateson Centre, University of Sheffield, Western Bank, Sheffield, S10 2TN, UK

**Keywords:** Heart morphogenesis, morphometry, extracellular matrix, 3D segmentation

## Abstract

Heart development involves the complex structural remodelling of a linear heart tube into an asymmetrically looped and ballooned organ. Previous studies have associated regional expansion of extracellular matrix (ECM) space with tissue morphogenesis during development. We have developed *morphoHeart*, an 3D image tissue segmentation and morphometry software which delivers the first integrated 3D visualisation and multiparametric analysis of both heart and ECM morphology in live embryos. *morphoHeart* reveals that the ECM undergoes regional dynamic expansion and reduction during cardiac development, concomitant with chamber-specific morphological maturation. We use *morphoHeart* to demonstrate that regionalised ECM expansion driven by the ECM crosslinker Hapln1a promotes atrial lumen expansion during heart development. Finally, we have developed a GUI that allows the morphometric analysis tools of *morphoHeart* to be applied to *z*-stack images of any fluorescently-labelled tissue.

## Introduction

Tissue morphogenesis in development requires the elaboration of simple structures into complex shapes. This includes common processes such as epithelial folding and tubular morphogenesis, requiring coordinated growth and shaping of multiple tissue layers or cell types. The developing heart is an excellent example of such a morphogenetic transformation. The embryonic heart initially forms a linear tube that comprises an outer myocardial tube and inner endothelial lining (the endocardium). This linear tube undergoes a complex morphogenesis that includes bending and looping of the tube to align the segments of the heart, and regional ballooning of the tube to start forming the chambers^1^. This morphogenesis is vital for establishing the blueprint of the heart, and is followed by substantive organ growth and formation of structures such as valves and trabeculae to support function. Therefore defects in early heart morphogenesis can have profound impacts on later heart structure and function^2^.

The extracellular matrix (ECM) is a crucial signalling centre in tissue development including the heart^3–5^ where it provides biochemical and biomechanical cues to overlying cardiomyocytes and endocardial cells. During early heart morphogenesis the cardiac ECM is rich in the glycosaminoglycan Hyaluronic Acid (HA) and the proteoglycan (PG) Versican which both play conserved roles in heart morphogenesis^6–10^. Both HA and Versican can sequester water^11,12^, allowing them to swell the ECM, increasing volume and hydrostatic pressure^13^. Hydrostatic pressure is increasingly recognised as a driver of tissue morphogenesis^14^, and HA-mediated expansion of ECM volume is important in several developmental contexts, including epithelial projection formation in the ear^15,16^, and initiation of the atrioventricular valve^17^. Furthermore, our previous work indicated that the cardiac ECM is asymmetrically expanded prior to heart tube morphogenesis through regionalised expression of the HA and proteoglycan cross-linking protein Hapln1a, and that this asymmetric ECM expansion is required for atrial morphogenesis^7^. Thus, analysing ECM-space in conjunction with detailed morphometric descriptors of the adjacent tissue will be vital to helping us to gain a better understanding of these matrix-tissue relationships during development.

Tools to analyse early heart morphogenesis in detail are limited. Recent studies in mouse have performed detailed quantitative characterisation of fixed samples^18–21^. However, fixed tissue analyses can have limitations, for example, collapse or shrinkage of tissue due to fixation, and alterations to the ECM (by modifying hydration or crosslinking), which may hamper efforts to understand volumetric ECM dynamics in the context of cardiac morphogenesis. Where possible, live analyses would address these issues, but imaging embryos that normally develop in utero, such as mouse, is challenging, and has been limited to stage-restricted analysis of embryos in live explant culture^22–24^. Zebrafish represent an excellent model in which to analyse early stages of heart development in live embryos: the embryos are transparent and develop externally, and early morphogenesis of the heart tube, together with genetic pathways underlying development and disease are well-conserved^25^. Limited morphometric segmentation of the zebrafish heart has been described in fish^26^ through manual segmentation of a single tissue layer with a limited number of defined parameters for analysis. While more complex morphometric tissue analyses are becoming more widely adopted, these often use bespoke code, or software that are not able to handle the complexity of the heart, and are not able to extract information about the acellular space between tissue layers, such as volumetric reconstructions of the cardiac ECM.

We have taken advantage of the zebrafish model for analysing heart morphogenesis, and used it to develop a new open-access image analysis software, *morphoHeart*, which allows multiparametric morphometric analysis of the developing heart in live embryos, including segmentation of the cardiac ECM. The design of *morphoHeart’s* graphical user interface (GUI) expands its use beyond that of cardiac tissue, facilitating analysis of multiple tissue layers, including label-free negative space segmentation of tissue or fluid within layers and division of tissue into segments or regions of interest for more granular analysis. Here we use *morphoHeart* to reveal new insights into early cardiac morphogenesis in the developing zebrafish.

## Design

Developing organs such as the heart are complex tissues, and their detailed morphology can be challenging to interpret using 2D images. Quantitative analyses of cardiac morphology are still limited, and label-free visualisation of ECM volume is not currently possible. Understanding the embryonic origins of congenital malformations requires refined quantitative analyses of heart morphology. These are generally restricted to fixed tissue, and currently no software can identify and analyse all layers that contribute to the heart.

Therefore, the core design goal for *morphoHeart* was to generate a visualisation and analysis software able to generate 3D reconstructions of the myocardium, endocardium and cardiac ECM of developing hearts from fluorescently labelled images, facilitating multi-parametric morphometric analysis of the tissue layers and heart morphology throughout development. This also includes visualisation and quantitative analysis not only of geometrical and volumetric parameters, but also tissue thickness, cell size and tissue expansion. Finally, we also wished to visualise conserved morphological features between biological samples, which can be hampered by heterogeneous morphology between individuals, and thus developed a method to standardise and overlay replicates to visualise conserved biological features.

These design goals required the incorporation of multiple methodologies, with features that can be user-configured to make the software adaptable to different needs. For example, negative-space segmentation of the ECM required a contour-based approach to create a volumetric reconstruction. Analysis of tissue deformation required definition of a line running through the centre of a specific 3D surface, and thus we incorporated centreline-finding methodology typically used to segment vasculature. Multi-sample analyses required integrated use of centrelines, user-defined cutting planes, and automated reslicing to generate quantitative representations of an ‘average’ heart.

*morphoHeart* was designed to be user-friendly, independent of programming experience. While it was originally developed for analysis of cardiac tissue, its GUI is designed to be appropriate for analysis of any type of fluorescently-labelled sample in which an image can be segmented from contours - including analysis of single or multiple layers, extraction of negative space volumes, and multi-segment and region analyses.

## Results

### Image acquisition, preprocessing and morphoHeart reconstructions

To segment the three layers of the heart (myocardium, endocardium, and ECM), *Tg(myl7:lifeActGFP); Tg(fli1a:AC-TagRFP)* double transgenic zebrafish embryos were imaged, in which the myocardium and endocardium are labelled. The heart beat was temporarily arrested, and *z*-stacks encompassing the heart were acquired (Fig S1A-C). To remove noise artefacts, accentuate details and enhance tissue borders, *z*-stacks are then processed, filtered, and cropped, and imported into *morphoHeart* for tissue layer segmentation (Fig 1A). To extract the myocardium and endocardium, individual slices making up each channel go through a process of contour detection and selection, extracting the contours that delineate the tissue layer (Fig 1B,C). Selected contours are classified either as internal or external, depending on whether they outline the lumen or the external borders of the tissue, respectively. Classified internal and external contours are used to create filled binary masks, one containing just the filled lumen of the tissue of interest (henceforth ‘filled internal contour mask’, Fig 1F,G), and another containing both the tissue layer, and the filled lumen (henceforth ‘filled external contour mask’, Fig 1D,E). For each channel, external and internal contour masks are combined to obtain a final mask delineating just the tissue layer itself (Fig 1H,I). Together this represents a total contour library from which all *morphoHeart* operations can subsequently be performed (Fig 1J). The resulting binary masks can be used to create volumetric 3D reconstructions (hereafter meshes) of each tissue layer of the heart (Fig 1K), facilitating the extraction of 3D morphological readouts to characterise heart morphogenesis.

**Figure 1.**
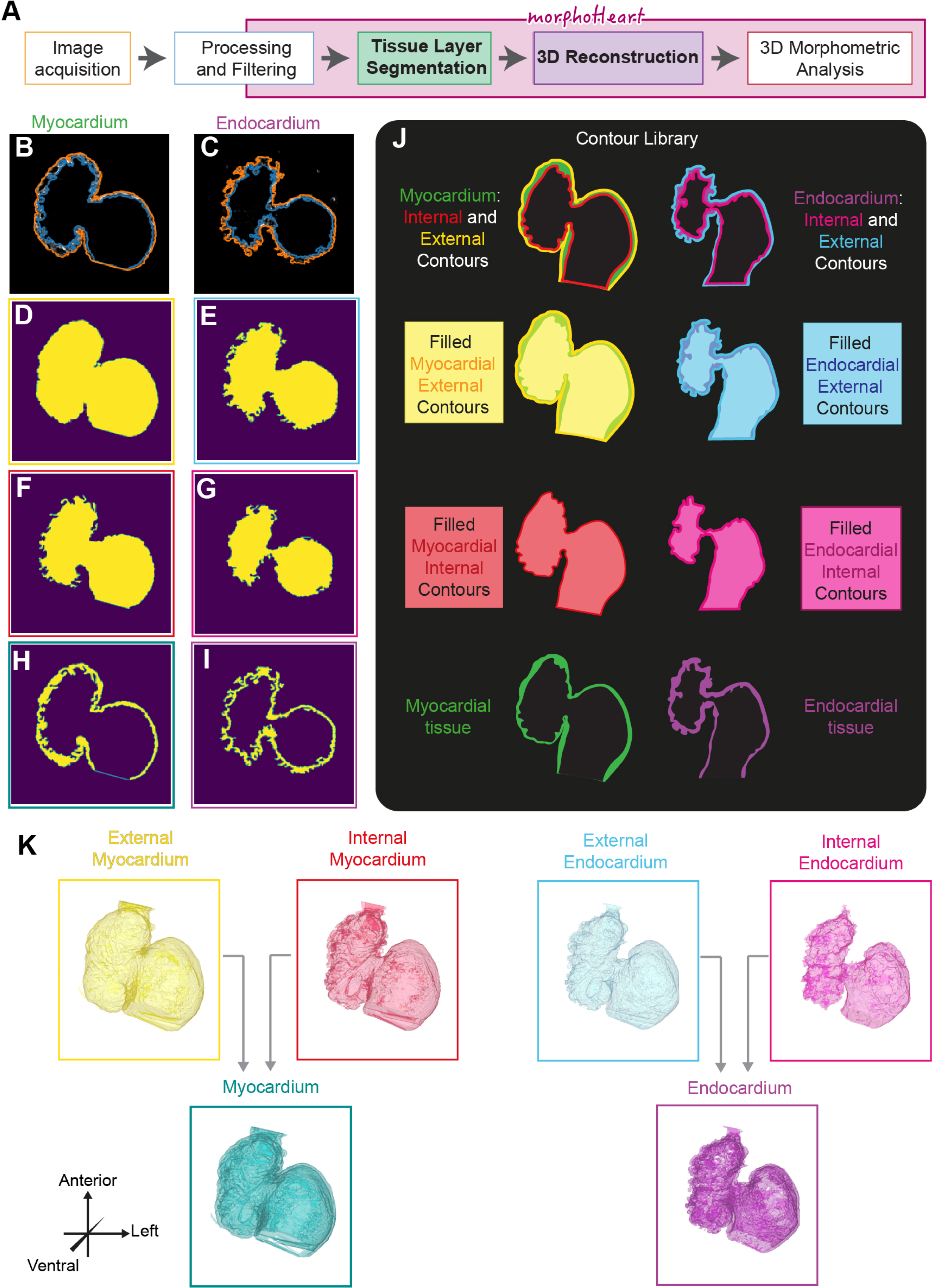
- morphoHeart image segmentation and volumetric tissue reconstructions. A: Schematic overview of the *morphoHeart* image processing and segmentation pipeline. B-I: Generation of tissue contour libraries. Identification of outer (orange) and inner (blue) contours of the myocardium and endocardium in single z-slices (B,C). For each slice masks are generated representing filled outer (D,E) and inner (F,G) tissue contours. Subtraction of inner from outer contours results in a tissue mask for each slice (H,I). J: Contour library generated by *morphoHeart*. K: 3D mesh reconstructions of external (yellow) and internal (red) myocardium, external (blue) and internal (pink) endocardium, and myocardial (green) and endocardial (magenta) tissues.

### The zebrafish heart undergoes periods of growth and compaction during early morphogenesis

To characterise the morphological changes the heart undergoes throughout looping and ballooning (Fig 2A-D), we imaged *Tg(myl7:lifeActGFP); Tg(fli1a:AC-TagRFP)* double transgenic zebrafish embryos at early looping (34hpf to 36hpf), after initial looping (between 48hpf and 50hpf), prior to onset of trabeculation (58hpf to 60hpf) and during early maturation of the chambers (72hpf to 74hpf), generating volumetric meshes of both the myocardium and endocardium (Fig 2E-H). Various methodologies have been previously described to quantify the extent of looping morphogenesis in zebrafish, relying on 2D image analysis^7,26–28^. Since the heart initially forms a linear tube, and looping is a morphological deviation from a linear state, we used the linear distance between poles and the length of the heart’s centreline to calculate a looping ratio, similar to our previous approach^7^, but taking into account the 3D organisation of the tissue. A heart ‘centreline’, (i.e. the line within the lumen of the heart, whose minimal distance in 3D from the tissue wall is maximal) (Fig 2I) was defined through calculation of a Voronoi diagram of the internal surface of the myocardium (Fig S2) and connected to the centre of the venous and arterial poles (which also serve as anchor points to calculate the direct linear distance between the poles). As looping proceeds, the linear distance between the poles reduces (Fig 2J), and between 34-50hpf the length of the centreline increases (Fig 2K), together resulting in an increase in the looping ratio (Fig 2L). Interestingly, while the linear distance reduces between 48-74hpf as the heart continues to compress between poles, the centreline length also decreases, resulting in maintenance of the looping ratio (Fig 2L). This supports the model that looping morphogenesis is completed by 2dpf and the chambers subsequently undergo other morphological rearrangements to shape the tissue.

**Figure 2.**
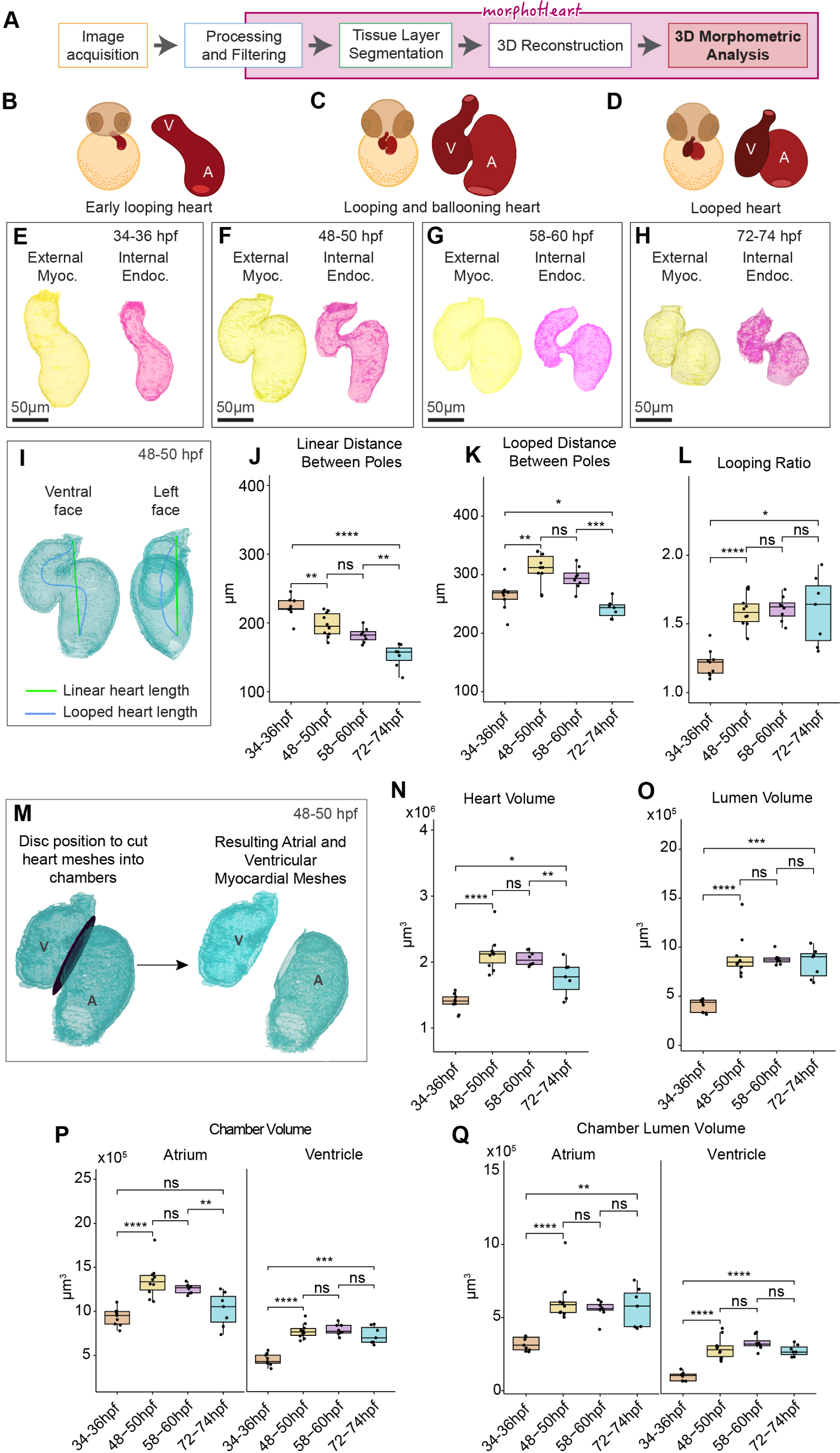
- The zebrafish heart grows and compacts during early cardiac morphogenesis. A: *morphoHeart* facilitates comprehensive 3D morphometric analysis. B-D: Schematic depicting the early stages of heart development analysed, including looping of the early tube (B, 34-36hpf), looping and ballooning (C, 48-60hpf), and the looped heart (D, 72-74hpf). E-H: Reconstructions of myocardial and endocardial meshes during heart development. I-L: Analysis of heart looping. Linear heart length (green line) and heart centreline (blue line) are extracted and measured (I). As the heart develops, the poles move closer together (J). During looping morphogenesis, the centreline’s looped distance elongates between 34-50hpf, and subsequently shortens (K). Looping ratio also increases between 34-50hpf, but then remains constant (L). M: Cardiac chambers can be separated via placement of a user-defined disc. N-O: Quantification of total heart volume reveals the heart increase in volume between 34-50hpf, and compacts again by 72-74hpf (N). Lumen volume increases with heart volume, and is then maintained (O). P-Q: Analysis of chamber volume reveals while both chambers grow between 34-50hpf, ventricle volume is maintained while the atrium shrinks (P). Lumen size in both chambers is maintained post-48hpf (Q). One-way ANOVA with multiple comparisons.* p<0.5, ** p<0.01, *** p<0.001, ns = not significant. 34-36hpf: n=9; 48-50hpf: n=10; 58-60hpf: n=8; 72-74hpf: n=7.

Ventral and lateral views of the heart suggest that chamber orientation changes during morphogenesis, in line with previously-described chamber realignments^27,29^. To characterise how distinct morphogenetic processes in individual chambers contribute to heart development, *morphoHeart* was developed with the functionality to divide meshes into sections through a user-defined plane, in this instance cutting through the atrioventricular canal and separating the chambers (Fig 2M). Isolation of the meshes for individual chambers, followed by definition of the pole and apex of each chamber, allows a more in-depth quantitative analysis of relative changes in chamber orientation during morphogenesis. The orientation of each chamber relative to the linear axis between the poles of the heart can be calculated from both a ventral view visualising how chambers align alongside each other (Fig S3 A-E), and from a lateral view visualising how chambers rotate relative to each other around the AVC (Fig S3 J-N). As the early heart tube undergoes looping, the ventricle pivots towards the right of the embryo or heart midline (Fig S3G-I), and the angle between both chambers reduces. This rearrangement may facilitate concomitant growth of both chambers and looping of the heart. After looping has finished, the ventricle has pivoted back to towards the left side of the embryo causing the angle between chambers to increase again. Analysis of chamber orientation from the lateral view demonstrates that again the position of the atrium remains relatively static, while the ventricle undergoes a rotation to initially become aligned more parallel with the linear plane of the heart, which is subsequently reversed post-looping (Fig S3O-R).

Visual inspection of the heart meshes suggests that the heart grows and shrinks between 34-72hpf (Fig 2E-H). Volumetric measurements of the external myocardium (as a proxy for whole heart size) and internal endocardium (as a proxy for lumen volume) were analysed. As the heart transitions from an early looping tube to a looped structure, the total volume of the whole organ, including the lumen, significantly increases (Fig 2N,O), expanding the blood-filling capacity of both chambers. Surprisingly, despite this growth, between 58-74hpf heart volume significantly reduces (Fig 2N). However, regardless of this reduction in heart volume, lumen capacity is maintained from 48hpf (Fig 2O), suggesting that remodelling of cardiac tissue occurs during early maturation, but this does not impact cardiac capacity. Individual analysis of chamber size reveals that both chambers grow between 34-50hpf (Fig 2P), accompanied by a substantial increase in lumen volume (Fig 2Q). However once the heart has undergone looping, the chambers display different dynamics, with the atrium reducing significantly in volume between 58-74hpf, while ventricular volume is maintained (Fig 2P). Similar to the whole heart analysis (Fig 2O), lumen size of both chambers is maintained (Fig 2Q), suggesting that the reduction in atrial volume is not due to a shrinkage of the whole chamber, but may instead represent a reduction in the amount of tissue.

### The cardiac chambers undergo distinct geometric changes during heart looping and chamber expansion

The atrial and ventricular chambers eventually adopt very different morphologies in the mature heart^30,31^. To understand the temporal geometric changes of the developing chambers, atrial and ventricular shape were quantified by measuring the dimensions of the ellipsoid that best fits the chamber myocardial mesh (Fig 3A, Fig S4A-F), including chamber width, length, depth, and asphericity (deviation of an ellipsoid from an sphere). During looping the atrium maintains its length and width, while increasing in depth (Fig S4G-I), suggesting this elongation in this axis is responsible for the significant increase in atrial size, potentially representing a ballooning-type growth. Meanwhile, the ventricle lengthens and narrows (Fig S4J-L), suggesting that enlargement of the ventricle during looping is due to chamber elongation. Once the heart has looped at 48-50hpf, the reduction in atrial volume we observed between 48-74hpf (Fig 2P) is accompanied by a gradual increase in depth and significant shortening of the atrium (Fig S4G-I), resulting in rapid adoption of a more spherical morphology by 72hpf (Fig 3B). In contrast to the atrium between 48-74hpf the ventricle only increases in depth (Fig S4J-L), maintaining its bean-shaped morphology (Fig 3C).

**Figure 3.**
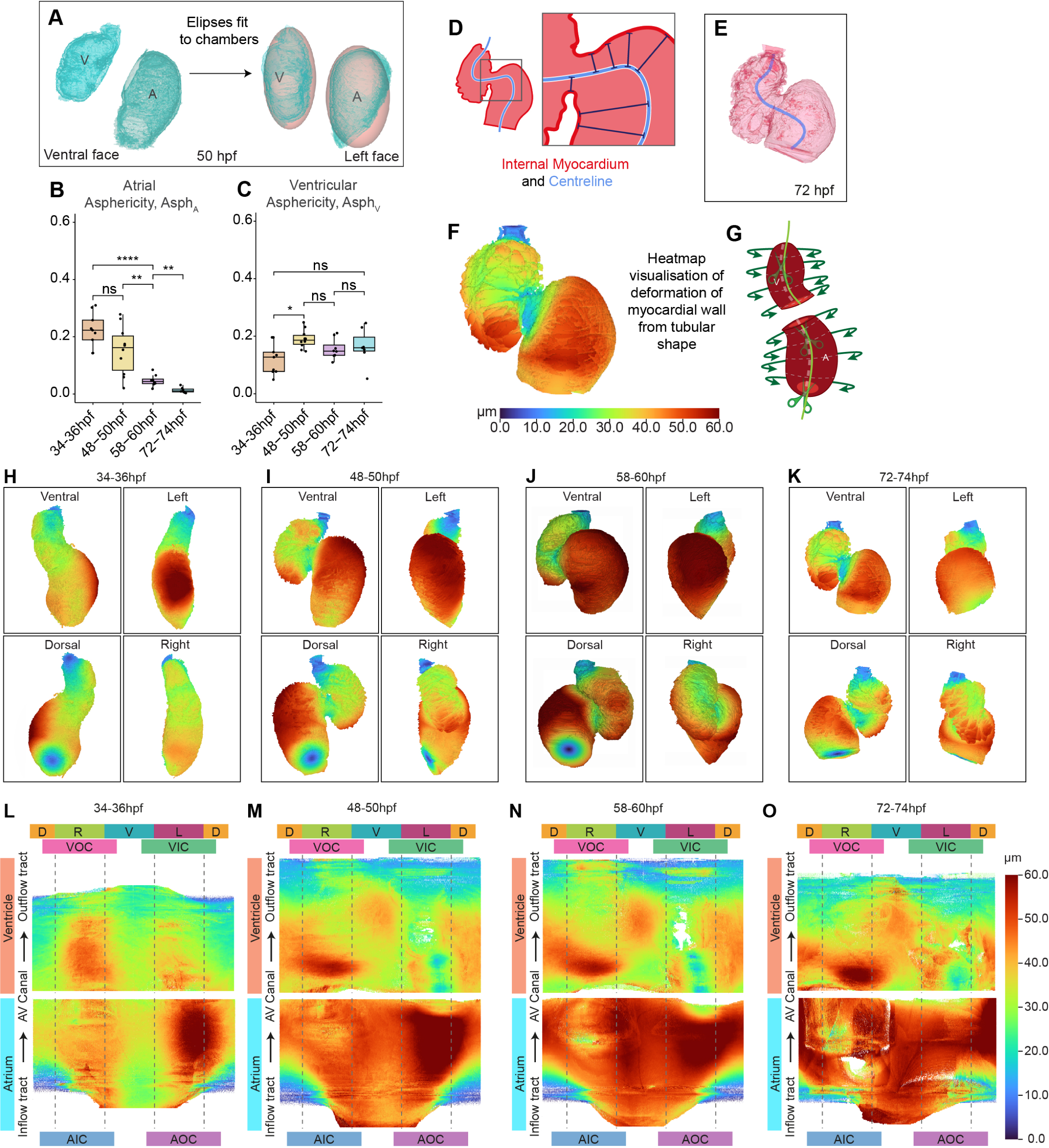
- Visualisation and quantification of chamber deformation reveals chamber-specific differences in growth. A-C: Ellipsoids are fitted to chambers to quantify chamber geometry (A). The atrium becomes more spherical (asphericity tends to 0) during development (B), while the ventricle initially becomes more aspherical by 50hpf with no further changes (C). One-way ANOVA with multiple comparisons.* p<0.5, ** p<0.01, *** p<0.001. D-G: Myocardial expansion/deformation can be quantified by measuring the distance between the myocardial centreline and the inner myocardial mesh (D). This value is mapped onto the inner myocardial mesh (E) using a heatmap to visualise 3D cardiac ballooning (F). 3D heatmaps can be unrolled into a standard 2D geometry for aggregation and comparison (G). H-J: Visualisation of 3D myocardial ballooning heatmaps identifies substantial deformation of the atrial outer curvature at 34-36hpf (H). By 48-50hpf this outer curvature deformation is enhanced, and the atrium is more ballooned than the ventricle. The ventricular apex can be seen emerging (I-K). L-O: Unrolled 2D ballooning heatmaps allows averaging of multiple hearts to identify conserved regions of deformation. By 74hpf deformation of the atrium has become more uniform (O). Labels around the 2D heatmaps indicate cardiac region: D - dorsal, V - ventral, L - left, R - right, AOC - atrial outer curvature, AIC - atrial inner curvature, VOC - ventricular outer curvature, VIC - ventricular inner curvature, AVC - atrioventricular canal. 34-36hpf: n=9; 48-50hpf: n=10; 58-60hpf: n=8; 72-74hpf: n=7.

Cardiac ballooning is a conserved process by which the chambers emerge from the linear heart tube^32,33^. In zebrafish, expansion of the chamber regions of the tube, a process akin to ballooning, has been described as occurring concomitant with cardiac looping from around 34hpf^34,35^, with the two processes together shaping the heart. To visualise and quantify regional expansion of the heart tube, the shorter distance between the heart’s centreline and the inner myocardial mesh was calculated throughout the heart (Fig 3D), and mapped onto the myocardial mesh using a colour-coded representation of chamber expansion (Fig 3F,H-K). This analysis reveals the emergence of the outer curvature of the atrium at 34hpf (Fig 3H) which becomes more pronounced as looping progresses (Fig 3I), when expansion of the ventricular outer curvature is also initiated. While analysis of individual 3D hearts is valuable for visualising localised regions of chamber expansion, it makes comparative analysis between biological replicates or stages relatively subjective. To enable the generation of an average heatmap of cardiac expansion (ballooning), combining multiple biological replicates per stage, *morphoHeart* uses the previously defined centreline to unroll each heatmap to a standard 2D matrix (Fig 3G, Fig S5A,B). Multiple samples with the same 2D format can then be combined to generate an average heatmap (Fig 3L-O, Fig S5C-D), which represents conserved geometry (Fig S5E-F), for example chamber expansion, from multiple replicates. Analysis of 2D ballooning heatmaps confirms that by 74hpf deformation of the atrium has become more uniform, representing the increase in sphericity we previously identified (Fig 3B), while the localised expansion of the ventricular apex becomes gradually more pronounced (Fig 3L-O).

### Individual cardiac chambers undergo separate processes of tissue growth and regional shrinkage

*morphoHeart* analyses of heart development highlight that cardiac chambers undergo geometric changes commensurate with ballooning; however, chamber size dynamics after initial looping indicate chamber expansion may not be a process only of tissue growth. We therefore investigated cardiac tissue volume in more detail (Fig 4A-F). Total myocardial volume increases as the heart undergoes looping morphogenesis, but surprisingly reduces again between 48-60hpf (Fig 4B). Analysis of the tissue volume of individual chambers reveals distinct chamber-specific myocardial tissue dynamics. Atrial myocardial tissue volume remains relatively consistent as the heart undergoes initial looping, but significantly reduces as the heart matures (Fig 4C). Conversely, ventricular myocardial volume increases significantly as the heart undergoes looping morphogenesis (Fig 4C), in line with the addition of second heart field cells to the arterial pole between 24hpf and 48hpf^36,37^, and subsequently remains constant. Analysis of endocardial tissue dynamics reveals a decrease in endocardial volume in the whole heart between 58-74hpf (Fig 4E) in both chambers (Fig 4F), which may reflect the general reduction in cardiac size at the later stage.

**Figure 4.**
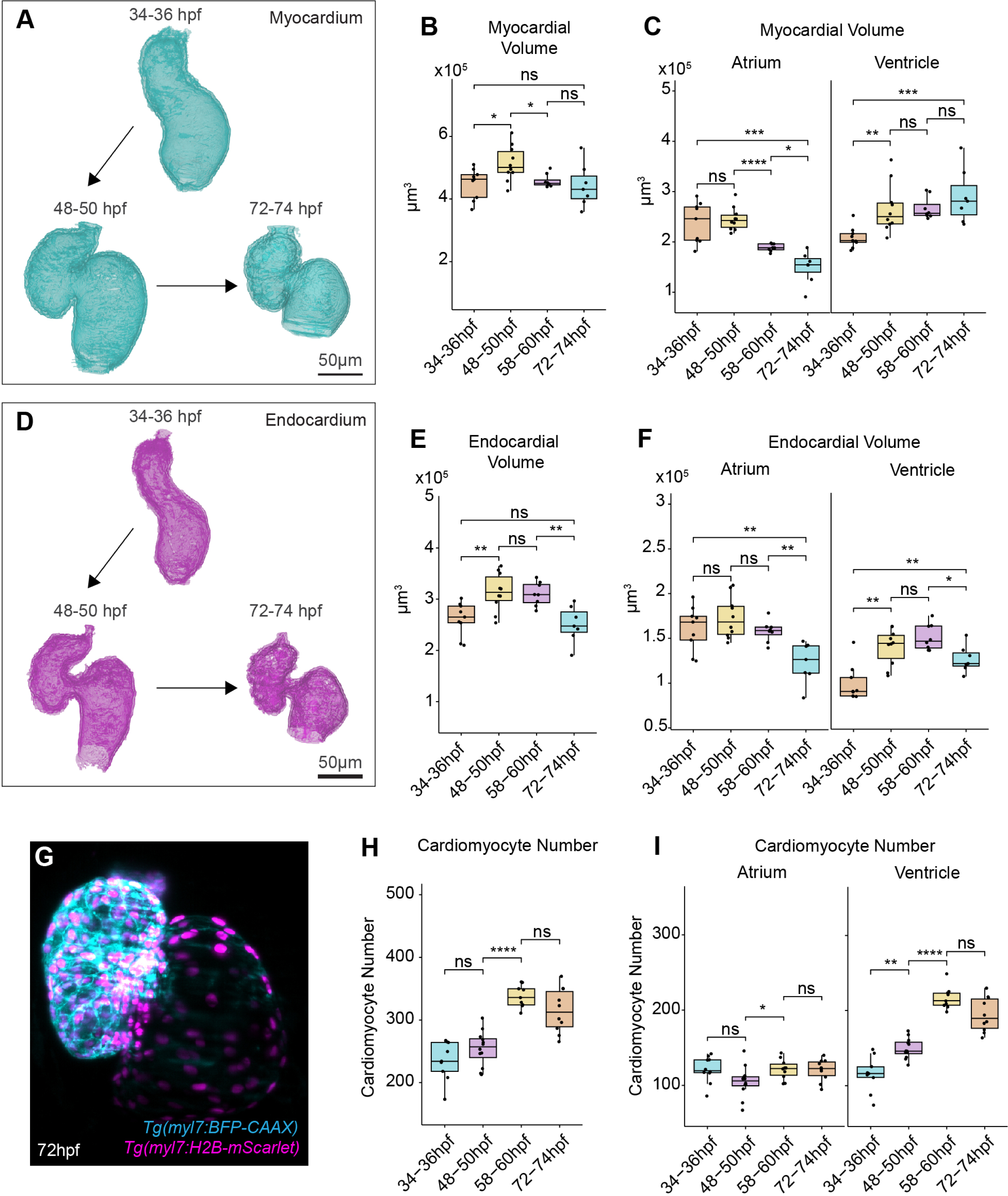
- The atrium and ventricle exhibit different tissue dynamics during morphogenesis. A-C : Quantification of myocardial tissue volume from myocardial meshes (A). Total myocardial volume increases during looping, and gets reduced at early stages of maturation (B). Chamber-specific analysis reveals a later reduction in atrial myocardium compared with an earlier increase and maintenance in ventricular myocardial volume (C). D-F: Quantification of endocardial tissue volume from endocardial meshes (D). Total endocardial volume decreases after 58hpf (E), driven by a reduction in endocardial tissue in both the atrium and ventricle (F). G-I: Quantification of cardiomyocyte number, from live lightsheet *z*-stack images of *Tg(myl7:BFP-CAAX);Tg(myl7:H2B-mScarlet)* (G). The total number of cardiomyocytes increases between 48-60hpf (H). Atrial cardiomyocyte number remains mostly constant (H), while ventricular cardiomyocyte number increases (I). One-way ANOVA with multiple comparisons.* p<0.5, ** p<0.01, *** p<0.001, **** p<0.0001, ns = not significant. 34-36hpf: n=8; 48-50hpf: n=10; 58-60hpf: n=7; 72-74hpf: n=7.

This reduction in atrial tissue volume after looping morphogenesis is complete at 48hpf is in line with our observation that total heart volume decreases over the same time frame driven primarily by a reduction in atrial size while lumen size is maintained (Fig 2). These changes in tissue volume could result from a reduction in number of cells, or a reduction in cell size. To address both these questions, we developed *morphoCell*, an integrated module in *morphoHeart* to perform cell analysis. We imaged *Tg(myl7:BFP-CAAX);Tg(myl7:H2B-mScarlet)* double transgenic embryos in which cardiomyocyte nuclei are labelled (Fig 4G), and used Imaris software to extract nuclei coordinates. These together with the myocardial *z*-stack were given as input into *morphoCell* where a plane was defined to separate chambers and allocate nuclei as atrial or ventricular (Fig S6B-D). This demonstrated that total cardiomyocyte number in the heart increases between 50-60hpf (Fig 4H), driven by the ventricle (Fig 4I), likely through continued differentiation of SHF cells^37^. However, cell number in the atrium remained constant, suggesting the reduction in atrial myocardial volume is not driven by cell loss or cell death, and together with previous studies ^36,38,39^ suggests that SHF addition to the venous pole predominantly occurs prior to 32hpf.

We therefore investigated whether atrial cardiomyocyte size reduces after initial heart looping morphogenesis. *morphoCell* can assign cardiomyocyte nuclei clusters, and measure 3D cardiomyocyte internuclear distance as a proxy for cell size (IND, Fig S6C-E). Analysis of total chamber cardiomyocytes revealed that atrial cardiomyocytes slightly increase in size during looping, but once looping has occurred only ventricular cardiomyocytes reduce in size (Fig 5A). Early chamber morphogenesis involves regionalised and chamber-specific changes in tissue morphology ^35^, and therefore we wished to assess cardiomyocyte size in more detail. Chamber-specific nuclei can be assigned to discrete regions of the chamber, such as the inner or outer curvatures, or dorsal or ventral face (Fig 5B). This revealed regional differences in atrial cardiomyocyte dynamics, with ventral and outer curvature cardiomyocytes associated with atrial expansion, and ventral atrial cardiomyocytes specifically reducing in size after looping (Fig 5C). Similarly, ventricular cardiomyocytes exhibit regional differences in expansion and reduction, with inner curvature cardiomyocyte size remaining relatively stable, while cardiomyocytes on the ventral, outer and dorsal faces of the ventricle undergo more dynamic changes in size (Fig 5D).

**Figure 5.**
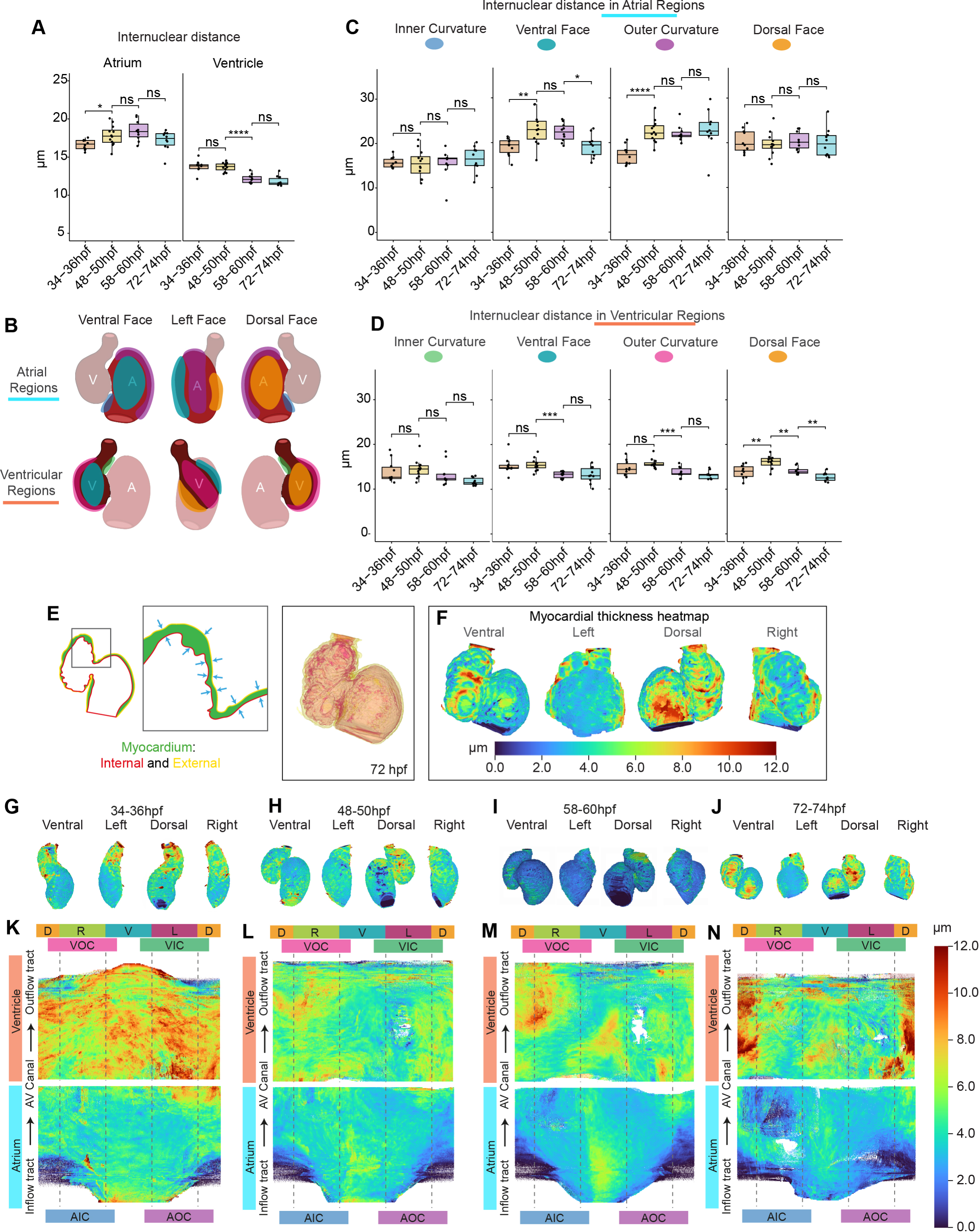
- Cardiac chambers undergo regionalised reduction in cell size. A-D : Quantification of internuclear cardiomyocyte distance as a proxy for cell size reveals an early increase in atrial cardiomyocyte size and a later reduction in ventricular cardiomyocyte size. Each chamber is subdivided into regions for more granular analysis (B). Growth and decrease in atrial cardiomyocyte size occurs predominantly in ventral and outer curvatures (C). Ventricular dorsal cardiomyocytes expand early, and all ventricular cardiomyocytes apart from those on the inner curvature subsequently decrease in size (D). Each dot represents the average internuclear distance, per region, in one heart. One-way ANOVA with multiple comparisons.* p<0.5, ** p<0.01, *** p<0.001. 34-36hpf: n=10; 48-50hpf: n=12; 58-60hpf: n=10; 72-74hpf: n=10. E-N: Myocardial wall thickness is quantified by measuring the distance between the inner and outer myocardial meshes (E), and mapped onto the outer myocardial mesh using a heatmap to visualise myocardial thickness in 3D (F). 3D myocardial thickness heatmaps (G-J) are unrolled to 2D (K-J), illustrating that the atrial wall is consistently thinner than the ventricular, and that both chamber walls thin during development. Labels around the outside indicate cardiac region: D - dorsal, V - ventral, L - left, R - right, AOC - atrial outer curvature, AIC - atrial inner curvature, VOC - ventricular outer curvature, VIC - ventricular inner curvature, AVC - atrioventricular canal.

Reduction in internuclear distance may reflect a change in cell geometry rather than a reduction in cell volume (i.e. cells get taller and narrower). Similarly, the decrease in atrial myocardial volume observed (Fig 4) is unlikely to be only attributed to a relatively modest and regional reduction in atrial cardiomyocyte size. We therefore sought to visualise myocardial wall thickness during development. As each tissue mesh comprises an outer and inner mesh, the shorter distance between these two meshes can be measured across the myocardial tissue (Fig 5E) and mapped onto the myocardial mesh as a heatmap (Fig 5F), providing a visual readout of myocardial thickness across development (Fig 5G-J). Analysis of 2D unrolled and averaged heatmaps demonstrates that the ventricular myocardial wall is consistently thicker than the atrial wall (Fig 5K-N). Importantly the atrial wall thins over development, which together with the regional reduction in cardiomyocyte IND supports the hypothesis that cardiomyocytes shrink after initial looping morphogenesis. Together this suggests that the chamber-specific changes in geometry that occur post-looping may be driven by regionalised changes in cell volume.

Our *morphoHeart* volumetric analysis thus suggests that concomitant with heart looping, the heart grows significantly through increase in cardiomyocyte size, accrual of cardiomyocytes and expansion of both chamber lumens until around 50hpf. Subsequently, cardiomyocytes undergo chamber-specific regional shrinkage while the lumen of the tissue is maintained, facilitating geometric changes that result in the adoption of specific ballooned morphologies in each chamber while maintaining cardiac capacity.

### The cardiac ECM undergoes dynamic regionalised and chamber-specific volumetric remodelling

We have previously shown that the cardiac ECM is regionalised prior to the onset of heart looping, where the atrial ECM is thicker than the ventricular ECM, and the left atrial ECM is expanded compared with the right^7^. We wished to investigate whether this regionalisation of ECM is maintained throughout early heart development, and how it relates to cardiac morphogenesis. We aimed to perform this analysis in live embryos (avoiding alteration of tissue morphology or matrix composition that may be introduced through fixation, dehydration, or processing), and without the use of ECM sensors (such as the previously-published HA sensor^6,41^), to avoid assumptions of ECM content. We took advantage of the contour libraries generated by *morphoHeart* to segment the negative space between the internal myocardial contour and external endocardial contour (Fig 6A), generating a mesh representing the cardiac ECM. Visualisation of cardiac ECM meshes across cardiac development (Fig 6B) revealed some expected features, such as a patchy reduction of the ECM in the ventricle between 48-74hpf, in line with previous reports in both mouse and zebrafish that ECM degradation occurs at the onset of ventricular trabeculation^42,43^.

**Figure 6.**
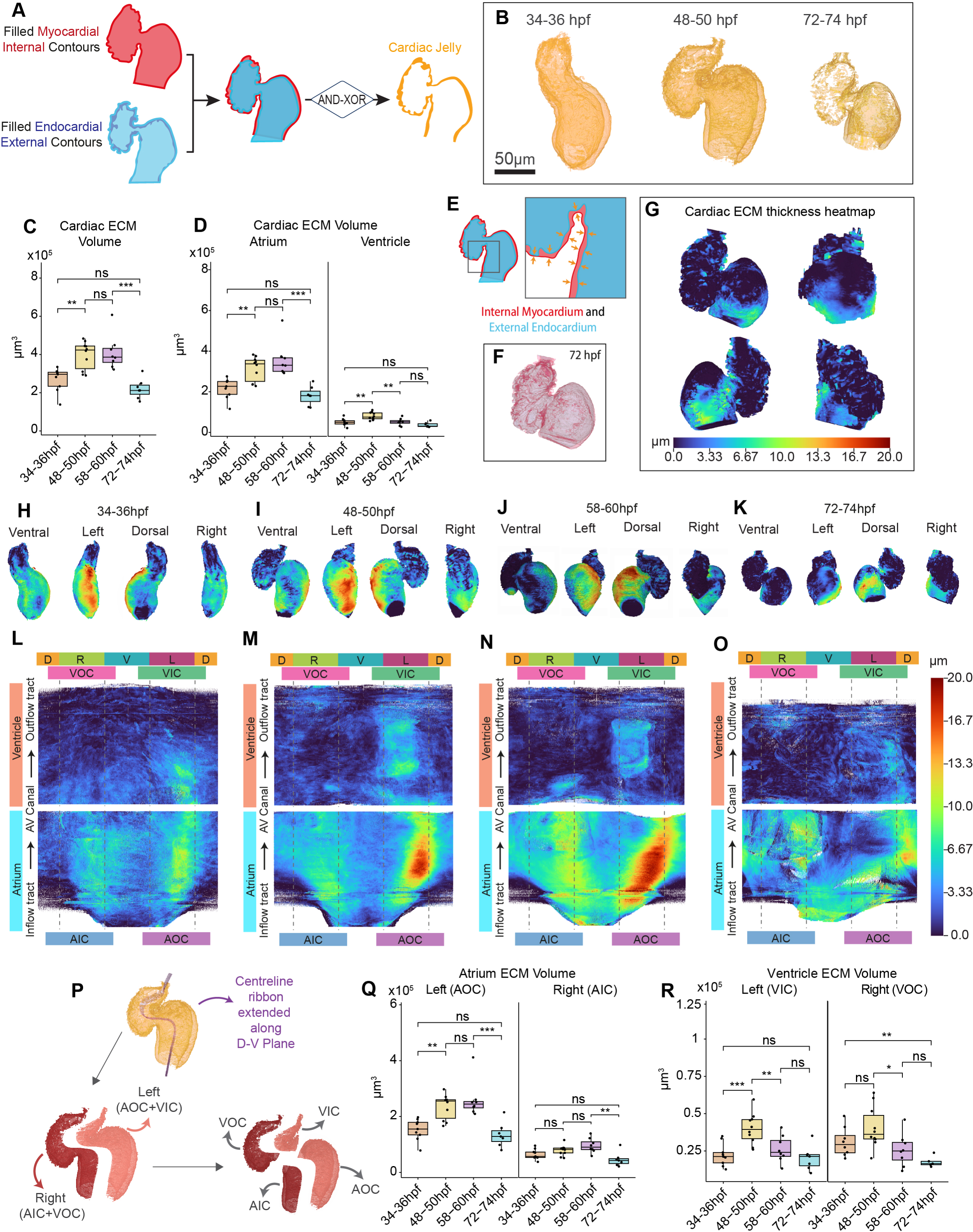
- The ECM undergoes chamber-specific regionalised expansion and reduction during heart morphogenesis. A: Schematic depicting the approach used to generate cardiac ECM meshes, by subtracting the filled external endocardial contour from the filled internal myocardial contour. B-D: Volumetric 3D reconstructions of the cardiac ECM during heart development (B), showing an apparent reduction in the ventricular ECM at 72-74hpf. Quantification of total cardiac ECM volume reveals a significant increase in ECM volume between 34hpf and 50hpf, followed by a reduction between 58hpf and 74hpf (C). The majority of cardiac ECM is found in the atrium (D), and while both chambers expand their ECM during looping, ventricular ECM reduces first between 48hpf and 60hpf, while atrial ECM is reduced only after 58hpf. ECM thickness is quantified by measuring the distance between the outer endocardial mesh and inner myocardial mesh (E), and mapped onto the inner myocardial mesh (F) using a heatmap to visualise ECM thickness in 3D (G). 3D heatmaps reveals the cardiac ECM is thicker in specific regions of the heart (H-K). Unrolled 2D ECM thickness heatmaps reveals the ECM is thicker in the atrium than the ventricle, and particular in the outer curvature of the atrium at 34-60hpf (L-N). The atrial ECM is still regionalised at 72-74hpf, but the thickening is repositioned to the dorsal face of the atrium (O). Labels around the outside indicate cardiac region: D - dorsal, V - ventral, L - left, R - right, AOC - atrial outer curvature, AIC - atrial inner curvature, VOC - ventricular outer curvature, VIC - ventricular inner curvature, AVC - atrioventricular canal. P: Schematic illustrating the cutting of the ECM mesh into left and right regions of both the atrium and ventricle. Q-R: Quantification of ECM volume in outer and inner curvatures of the atrium (Q) and ventricle (R) reveal the regionalised dynamics that drive cardiac ECM expansion and reduction. One-way ANOVA with multiple comparisons.* p<0.5, ** p<0.01, *** p<0.001, ns = not significant. 34-36hpf: n=9; 48-50hpf: n=10; 58-60hpf: n=8; 72-74hpf: n=7.

Quantitative analysis revealed that the ECM is highly dynamic, first significantly expanding between 34-50hpf as the heart loops, before reducing between 58-74hpf as the heart matures (Fig 6C). Chamber-specific analysis revealed that atrial ECM volume is consistently higher than ventricular ECM volume, and while both chambers exhibit the same types of dynamics, the timing is different, with the ventricular ECM reducing between 48-60hpf, earlier than the atrial ECM which reduces only between 58-74hpf (Fig 6D). This suggests that the chambers have distinct mechanisms for managing ECM degradation or reduction.

To visualise ECM thickness, we used *morphoHeart* to measure the distance between the ECM mesh contours (Fig 6E) and mapped the thickness values onto the external ECM tissue contour, using a heatmap scale to visualise ECM thickness (Fig 6G). Inspection of 3D ECM heatmaps throughout development reveals that the cardiac ECM is highly regionalised, with an expansion of the ECM on the outer curvature of the atrium at 34-60hpf (Fig 6H-J), which is repositioned to the dorsal atrial face by 74hpf (Fig 6K). 2D unrolled and averaged heatmaps facilitated a granular analysis of ECM regionalisation. At 34hpf the ECM is thickest on the left (outer) curvature of the atrium, in line with our previous findings at 26hpf^7^. We also observed a mild localised ECM thickening on the right (inner) curvature of the atrium and the left-sided ECM thickening expands into the proximal ventricle (Fig 6L). At 48-60hpf this regionalised thickening of the ECM in the atrium is maintained, and the magnitude increased, on both outer and inner curvatures (Fig 6M-N), while the ECM thickening in the inner curvature of the ventricle is slightly reduced at 60hpf (Fig 6N). By 72hpf the atrial ECM is still regionally expanded, but the left-sided expansion is now positioned to the dorsal face (Fig 6O). The left-sided inner ventricular ECM expansion has reduced, although the ECM in that region still appears slightly thicker than the right side (outer curvature), which may be in line with regionalisation of trabeculation onset^43^. To quantify ECM volume specifically in these chamber regions, *morphoHeart* used the centreline to divide each chamber into left and right sides (Fig 6P). This confirmed that the atrial ECM is greater on the left side, and expands more significantly to amplify the magnitude of the asymmetry as the heart undergoes morphogenesis (Fig 6Q). While left and right ventricular ECMs are more similar in volume, the left side undergoes a more dynamic expansion and reduction (Fig 6R), in line with the changes in thickness depicted in the heatmaps. *morphoHeart* therefore reveals novel chamber-specific dynamics in ECM expansion and reduction during cardiac morphogenesis.

### ECM crosslinker Hapln1a promotes cardiac growth dynamics

We previously demonstrated that the ECM crosslinker Hapln1a is required for regulating early ECM volume asymmetries and heart growth^7^. To validate that *morphoHeart* can deliver more detailed analyses of mutant phenotypes we performed *morphoHeart* analysis of *hapln1a* mutants at 34-36hpf, 48-50hpf and 72-74hpf (Fig 7A,B). Analysis of heart size revealed that *hapln1a* mutant hearts are only significantly smaller than siblings at 48hpf (Fig 7C), once the heart has undergone morphogenesis. This reduction in heart size is driven by a failure of the *hapln1a* mutant atrium to expand by 48hpf (Fig 7D,E). Analysis of both lumen and myocardial volume reveals that defective atrial growth is driven by limited expansion of the *hapn1a* mutant lumen (Fig 7F-J), demonstrating that Hapln1a links atrial ballooning to lumen expansion.

**Figure 7.**
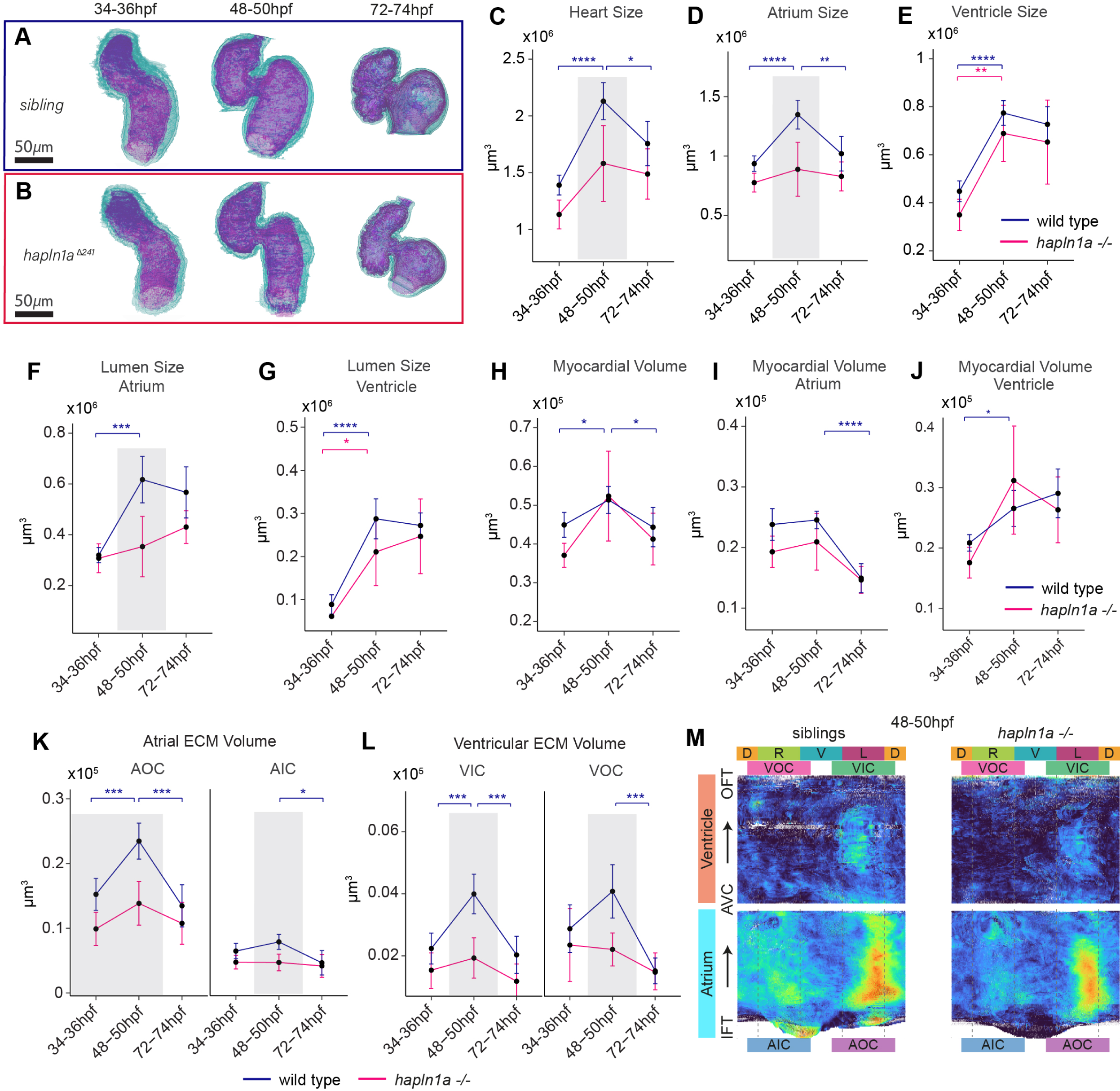
- hapln1a mutants exhibit defects in myocardial dynamics and ECM expansion. A-B: Myocardial (green) and endocardial (magenta) 3D reconstructions of sibling (A) and *hapln1a* mutant hearts (B) at 34-36hpf, 48-50hpf and 72-74hpf. C-E: Quantification of heart size reveals that *hapln1a* mutant hearts (pink) are smaller at 34-36hpf and 48-50hpf than wild type siblings (blue, C), largely due to a reduction in atrial size during early looping (D,E), and a failure of the atrium to balloon by 48-50hpf (D). F-J: Quantification of lumen size and myocardial tissue volume shows that *hapln1a* mutants fail to expand the atrial lumen at 48-50hpf (F), whereas the ventricular lumen is unaffected (G). K-M: Analysis of regional ECM volume in wild type siblings and *hapln1a* mutants. ECM volume doesn’t expand in either the atrium or ventricle of *hapn1a* mutants at 48-50hpf compared to wild type siblings (K,L). 2D ECM thickness heatmap reveals that while the magnitude of ECM expansion and regionalisation is reduced in *hapln1a* mutants, a small region of the atrium still exhibits a thicker ECM (M). Asterisks indicate significant difference between time points (blue indicates significance in siblings, pink indicates significance in *hapln1a* mutants). Grey boxes indicates significant difference between wild type siblings and mutant at the indicated timepoint. Two-way ANOVA with multiple comparisons.* p<0.5, ** p 0.01, p<0.001, ns = not significant. Labels around the heatmap indicate cardiac region: D - dorsal, V - ventral, L - left, R - right, AOC - atrial outer curvature, AIC - atrial inner curvature, VOC - ventricular outer curvature, VIC - ventricular inner curvature, AVC - atrioventricular canal.

Finally, we examined ECM volume and distribution in *hapln1a* mutants. The overall volume of cardiac ECM is reduced (Fig 7K,L), resulting in a diminished contribution of the cardiac ECM to the total heart volume (Fig S7A-C). We observed that *hapln1a* promotes the regionalised expansion of ECM between 34-50hpf in both the atrium and ventricle (Fig 7K,L), as well as apparently protecting the ECM at the inner curvature of the ventricle from premature degradation by 74hpf (Fig 7L). Surprisingly, despite this significant reduction in regional ECM volume in the *hapln1a* mutant, averaged 2D ECM thickness heatmap analysis of *hapln1a* mutants at 48hpf reveals a small area of the atrial outer curvature retains some expansion (Fig 7M), although the magnitude and expanse of this thickening appears reduced compared to wild type, in line with quantitative analysis (Fig 7K).

Together this suggests that *hapln1a* plays a broader role in amplifying the magnitude of asymmetries within the cardiac ECM, and that other genes are likely acting together with *hapln1a* to generate asymmetries within the cardiac ECM to help promote atrial expansion during cardiac morphogenesis.

## Discussion

### morphoHeart reveals new insights into cardiac morphogenesis

Early heart morphogenesis is a complex asymmetric process that requires the timely coordination of distinct events, including looping and regional ballooning of the linear heart tube. Using *morphoHeart*, we have demonstrated that the complex 3D morphological transformations of the zebrafish heart tube during cardiac development can be characterised through comparative integrated analysis of 3D morphometric parameters in wild-type hearts at key developmental stages.

*morphoHeart’s* quantitative results of myocardial growth during early looping shows that the increase in myocardial mass is driven by growth of the ventricle, while the atrial myocardium remains constant, corroborating previous studies showing that second heart field addition occurs earlier at the venous pole than the arterial^36^, likely prior to the stages we capture here. We further show for the first time an atrial-specific reduction in total myocardial volume after initial looping morphogenesis, while ventricular myocardial mass is maintained. *morphoHeart’s* capability to perform integrated analyses demonstrate that this reduction in atrial myocardium, and maintenance in ventricular myocardium are both associated with regionalised reduction in cardiomyocyte size, but in the ventricle myocardial volume is maintained through increased cell numbers. Chamber-specific analysis highlighted that these differences may be the result of ongoing chamber-specific refinement mechanisms; for example, the increase in ventricular cardiomyocytes could be due to ongoing addition of cells to the arterial pole from the SHF, and/or through the proliferation of cardiomyocytes during trabecular seeding^44^. Furthermore, the chamber-specific regional reductions in cell size we observe are in line with other studies that suggest that anisotropic cell shape changes drive tissue remodelling^27,40,45–47^.

Our data suggests that movement of the ventricle primarily drives heart looping. We observe a combination of frontal and sagittal rotations in the ventricle suggesting that, contrary to the findings of previous studies^27^, the deformation of the linear heart tube into a S-shaped loop does not solely take place in the frontal plane. Studies in other models have not only corroborated the three-dimensionality of looping morphogenesis process by describing the sequential frontal (left/right) and transverse (cranial/caudal) rotations of the chambers and OFT involved in looping morphogenesis^48–50^, but also described the principal role played by the ventricle during this asymmetric process of looping^51,52^. This suggests that the ventricular rotations underpinning chamber rearrangements during cardiac looping morphogenesis are conserved across species.

### The cardiac ECM undergoes regionalised dynamic changes in volume

*morphoHeart* allows the first 3D volumetric visualisation and analysis of the cardiac ECM in live embryonic hearts. We have shown that ECM expansion in both chambers is associated with the initial growth of the heart during looping and ballooning morphogenesis while ECM reduction, possibly driven by degradation, dehydration, or compaction, is subsequently linked to chamber-specific remodelling and maturation. Reduction in ECM volume occurs earlier in the ventricle than the atrium (between 48-50hpf compared to 58-74hpf), corresponding with the onset of ventricular trabeculation, which has been linked to specific dynamics of ECM remodelling^42,43,53^.

ECM thickness analysis provides a more granular understanding of cardiac ECM distribution, including regional expansion of the ECM on the left side of the heart tube, in line with our previous observations^7^ and previous studies describing the cardiac ECM of embryonic chick hearts as thicker on the left-right regions of the heart tube compared to the antero-posterior^54–56^. We also demonstrate for the first time that this regional expansion is maintained as the heart undergoes looping, chamber expansion, and early maturation, although in the atrium, the asymmetric expansion switches from the left face to the dorsal face.

Our analysis further revealed a distinct region of thickened ECM in the inner curvature of the atrium close to the venous pole, in the same area where previous studies have located the zebrafish pacemaker/sinoatrial node cells^36,57^. Studies in animal models have identified that the cells comprising the sinoatrial node are embedded within a biochemically and biomechanically distinct ECM that serves as a protective scaffold to pacemaker cardiomyocytes, reducing the mechanical strain and mechanotransduction they would experience from cardiac contractility^58^, raising the possibility that these cells are similarly isolated in the zebrafish heart during development.

We reveal a novel requirement for in driving expansion of the atrial lumen, drawing parallels with studies in Drosophila demonstrating that proteoglycans and glycoproteins regulate expansion of the intestinal lumen^60^, heart lumen^61^, and interrhabdomeral space in the eye^62^. The role of Hapln1a in regulating ECM and atrial expansion likely stems from its function in the ECM. Hapln1a is a cross-linking protein that mediates the interaction between HA and proteoglycans^63–65^, which in turn provides structure and biomechanical cues to tissues. Although proteoglycans and HA can interact and form complexes in the absence of link proteins, *in vitro* studies have shown that cross-linking allows the ECM to sustain higher loads (i.e. increased compressible resistance) whilst maintaining an elastic structure^65,66^. In addition, a stable and elaborate network of HA-PG complexes into which GAG chains can sequester water could result in formation of a hydrated and regionally expanded matrix that promotes atrial wall deformation, as well as signal transduction and proliferation through mechanical tension which has been proposed to correlate endocardial growth with myocardial ballooning^67^.

Importantly, our detailed 3D analysis of ECM thickness reveals that in *hapln1a* mutants, while ECM volume is significantly reduced at 48hpf, and the magnitude of matrix asymmetry is diminished, a small patch of thickened ECM remains in the outer curvature of the atrium. This demonstrates that while Hapln1a promotes the magnitude of ECM asymmetry, one or more additional components must also contribute. Whether this is due to regionalised production of HA, proteoglycans, or other ECM crosslinking proteins, or rather is due to localised expression of ECM-degrading enzymes, is unclear. However, the detailed insights provided by *morphoHeart* now highlight the complexity of dynamic ECM composition in shaping the developing heart.

### morphoHeart, a new tool for morphometric analysis

To date, a single tool or pipeline cannot address all the processing and analytic requirements for analysis of 3D datasets. Fully automatic 3D segmentation of biological images can be computationally demanding and can perform poorly due to low local contrasts, high noise levels and signal from structures or artefacts surrounding the objects of interest^68,69^. Once the structures of interest have been segmented, either by fully automatic or manual segmentation, limited 3D object quantification in open source software (e.g. 3D Viewer^70^, MorphoLibJ^71^, 3D Slicer^72^) extracts few quantitative readouts, restricting the depth of the analysis. Some commercially available software specifically designed to analyse medical (e.g. Mimics Materialise, Belgium) or biological (e.g. arivis Vision 4D, Germany; Imaris, Oxford Instruments; Volocity, PelkinElmer) images provide a more extensive toolset for the morphological analysis of 3D objects. Nevertheless, the available metrics might not meet all the user-specific needs. *morphoHeart* is an exciting alternative offering a comprehensive pipeline that provides semi-automated segmentation capabilities and improves the suite of tools and quantifications available in the biological field to characterise organ morphology. While *morphoHeart* was initially developed for the study of zebrafish heart development, it can be applied to studying a wide range of morphogenetic processes in other organs and organisms, contributing to our understanding of the tissue organisation mechanisms and morphogenesis processes underpinning them.

Previous studies have used either finite element (FE) models or 3D computational simulations to study the asymmetric process of heart tube morphogenesis^19,73^. Using simple representations of the heart tube and a combination of constraints and conditions, the models have recapitulated the bending and torsion of the heart, gaining insights into the internal and external forces involved in the formation of the mammalian helical loop. The morphological quantifications, geometries and 3D reconstructions provided by *morphoHeart* could be fed into similar descriptive or predictive models to better define the mechanisms, including the relationship between the cardiac tissue layers, that drive heart looping and chamber ballooning^19,73–76^.

In addition to the comprehensive morphometric quantifications performed by *morphoHeart*, it also provides the first negative space segmentation of the cardiac ECM. A previous study used automated constrained mesh inflation and subtraction to define the negative space surrounding joints^77^; however, our contour-directed negative space segmentation allows a more detailed analysis of regional expansions of the segmented matrix. While transgenic lines providing fluorescent readouts of Hyaluronic Acid have been described^6,41^ which could be used for analysis of the cardiac ECM in *morphoHeart*, these lines are not specific to the heart, making clean segmentation challenging. In particular, if ECM composition changes over time, the use of ECM sensor lines could render specific stages difficult to analyse, whereas *morphoHeart’s* segmentation approach of ECM volume from the negative space makes no assumptions about ECM content.

### Limitations

To the author’s knowledge, no other analytic approaches have been published quantifying the cardiac tissues of the developing zebrafish heart or chambers (including the cardiac ECM) with the resolution and detail provided by *morphoHeart*, making it challenging to compare and validate *morphoHeart’s* performance. However, we hope the morphometric characterisation of wild-type embryos delivered here can become the benchmark against which future studies can be compared.

*morphoHeart* offers a novel suite of morphometric parameters for performing detailed quantifications and characterisations of heart morphogenesis. Some of the processes used to generate data (for example tissue ballooning) requires the use of a centreline, which assumes a tubular nature to the structure that may not be appropriate or applicable in other tissue contexts. Thus, some functionality may remain limited to specific scenarios. However, the open-source nature of *morphoHeart* will allow other researchers to develop its analysis capabilities, implementing new quantifications or descriptors that support morphometric analysis of other tissues.

The processing and filtering steps used prior to *morphoHeart* segmentation were optimized for the myocardial and endothelial markers in the *Tg(myl7:lifeActGFP); Tg(fli1a:AC-TagRFP)* double transgenic line. These steps will likely require optimization dependent upon the transgenic line and image quality, to allow accurate contour demarcation and tissue layer segmentation.

Finally, due to the approach used to obtain undisturbed 3D image datasets of the whole heart (temporary cessation of heart beat) hearts were analysed with both chambers in a ‘relaxed’ state, which is not directly representative of any stage in the cardiac cycle. Despite this limitation, the same approach was used for all the analysed embryos and so the morphometric parameters obtained at the multiple developmental stages are comparable. The use of adaptive prospective optical gating^78^ or macroscopic-phase stamping^79^ imaging techniques in future studies will allow the acquisition of datasets at specific phases or throughout the whole cardiac cycle, which if combined with *morphoHeart’s* capabilities could provide deeper understanding of the dynamic changes in tissue morphology during cardiac contraction.

*morphoHeart*, along with a detailed user manual, are available for download from https://github.com/jsanchez679/morphoHeart.

## Methods

### Resources

| REAGENT OR RESOURCE | SOURCE | IDENTIFIER |
| --- | --- | --- |
| **Experimental Models (organisms/strains)** |  |  |
| *Tg(myl7:LifeActGFP)* | See Reishauer et al.<sup>80</sup> | s974Tg |
| *Tg(myl7:BFP-CAAX)* | See Guerra et al.<sup>81</sup> | bns193Tg |
| *Tg(myl7:H2B-mScarlet)* | See Boezio et al.<sup>82</sup> | N/A |
| *Tg(fli1:Actin-CB-TAGRFP)* | See Savage et al.<sup>83</sup> | sh511Tg |
| *hapln1a<sup>sh580</sup>* | See Derrick et al.<sup>7</sup> | sh580 |
| **Software and algorithms** |  |  |
| Vision4D | arivis | https://www.arivis.com/ |
| *morphoHeart* | This study | https://github.com/jsanchez679/morphoHeart |
| FIJI | See Schindelin et al.<sup>84</sup> | https://fiji.sc/ |
| Pre-*morphoHeart* Masking and Cropping FIJI Macro | This study | See Supplemental Data |
| **Chemicals, Peptides, and Recombinant Proteins** |  |  |
| Ethyl 3-aminobenzoate methanesulfonate salt (Tricaine) | Sigma | A5040 |
| 1-phenyl2-thiourea (PTU) | Sigma | P7629 |

### Zebrafish husbandry

Adult zebrafish were maintained according to standard laboratory conditions at 28.5°C. Embryos older than 24hpf were treated with 0.2 mM 1-phenyl-2-thiourea (PTU) in E3 medium to inhibit melanin production. All animals were euthanized by immersion in overdose of Tricaine (1.33g/l). Animal work was approved by the local Animal Welfare and Ethical Review Body (AWERB) at the University of Sheffield, conducted in accordance with UK Home Office Regulations under PPL PA1C7120E, and in line with the guidelines from Directive 2010/63/EU of the European Parliament on the protection of animals used for scientific purposes.

### Lightsheet imaging

To assess cardiac morphology at different developmental stages, live or fixed zebrafish embryos were imaged on a ZEISS Lightsheet Z.1 microscope. To stop the heartbeat of live embryos and aid image analysis, prior to mounting, 3 to 5 embryos were anesthetised by transferring them from a dish containing E3+PTU to a new cooled dish containing E3 and 8.4% Tricaine (E3+Tricaine). Anesthetised embryos were embedded in 1% low melting point agarose with 8.4% of Tricaine in black capillaries (1mm diameter; Brand 701904). To ensure the heartbeat was arrested during the acquisition, the imaging chamber was filled with E3+Tricaine and maintained throughout the experiment at 10°C.

All images were acquired using a 20X objective lens with 1.0 zoom. Single-side lasers with activated pivot scan were used for sample illumination. High-resolution images capturing the whole heart were obtained with 16 bit image depth, 1200 x 1200 pixel (0.228µm x 0.228µm pixel size resolution) image size and 0.469-0.7µm *z*-stack interval. For double fluorescent transgenic embryos, each fluorophore was detected on separate channels.

### Image preprocessing

To remove noise artefacts, accentuate details and enhance tissue borders, raw lightsheet *z*-stacks for each tissue channel were processed and filtered in arivis Vision4D. To smooth noisey regions but preserve the edges of each tissue layer, the *Denoising Filter (3D)* was applied to the RAW dataset. The resulting images were then processed using the *Background Correction* filter to reduce variations in intensity throughout the whole image set. Next, the *Morphology Filter* was used to sharpen the tissue borders, followed by *Membrane Enhancement* to boost the signal of membranes, producing clear slices with enhanced and sharpened borders in each channel.

To further enhance signal and reduce file size, after individual channels were processed in Vison4D, images were then processed in Fiji. First any residual salt-and-pepper noise was removed using the *Despeckle* filter. An *Enhancement* filter was then applied to both channels to improve the contrast of the images without distorting the grey level intensities. Finally, a Maximum Intensity Projection (MIP) of a composite containing both processed channels was used to define a square that contains the region of interest (ROI) comprising the heart. This ROI was used to crop each channel reducing the image size to be imported into *morphoHeart* for segmentation.

### Statistical analysis

All data was analysed and plotted in R. Significant differences between stages were analysed using one-way ANOVA followed by a Tukey post-hoc test. Significant differences between stages and/or genotypes were analysed using two-way ANOVA followed by a Tukey post-hoc test.

## Acknowledgements

We are grateful to Eric Pollitt, Emma Armitage, Chris Chan Jin Jie, Yangsheng Zhou, Enze Wang and Angelica Spadaro for testing early versions of *morphoHeart*. We thank Chris Derrick, Eric Pollitt, and Tanya Whitfield for critical reading of the manuscript. Lightsheet imaging was performed at the Wolfson Light Microscopy Facility using Zeiss Z1 lightsheet microscopes (BBSRC ALERT14 award BB/M012522/1 and BHF Infrastructure Grant IG/15/1/31328). J.S-P is supported by BBSRC Standard Grant BB/W004305/1. E.N is supported by a British Heart Foundation Fellowship award FS/16/37/32347.

## Author Contributions

Conceptualisation, J.S-P and E.N; Methodology, J.S-P and E.N; Software, J.S-P; Investigation, J.S-P and E.N; Resources, J.S-P; Data Curation, J.S-P and E.N; Formal Analysis, J.S-P and E.N; Writing – Original Draft, J.S-P and E.N; Writing – Review & Editing, J.S.P and E.N; Visualisation, J.S-P and E.N; Funding Acquisition, E.N; Supervision, E.N.

