## Supplemental Data for "*morphoHeart*: a novel quantitative tool to perform integrated 3D morphometric analyses of heart and ECM morphology during embryonic development"

+ Lead author

**Supplemental Data:**
Supplemental Figures S1-S7
Pre-*morphoHeart* Masking and Cropping FIJI Macro

Figure S1

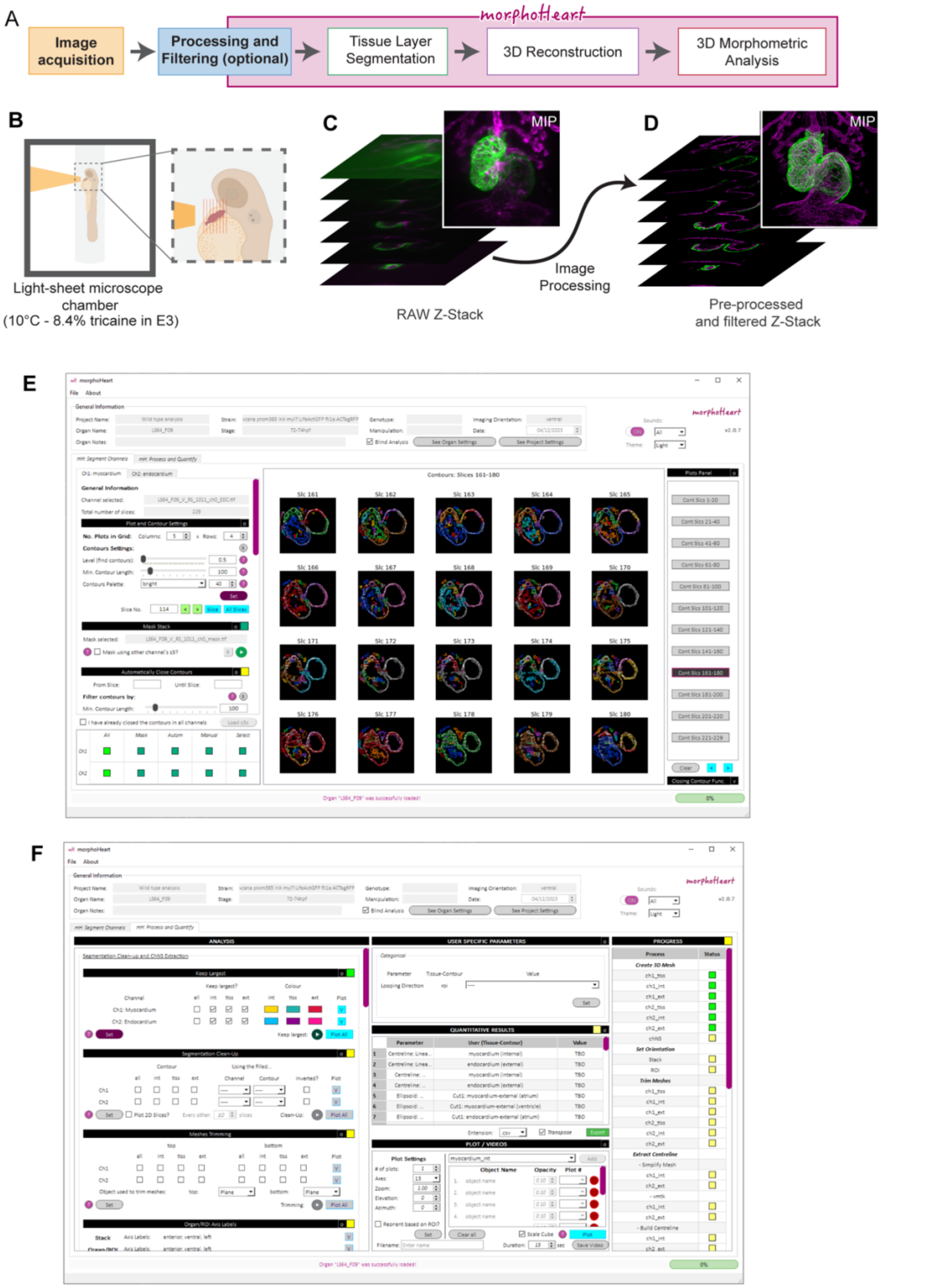


*Figure S1 - Image acquisition and the morphoHeart GUI.*

A: Schematic overview of the *morphoHeart* pipeline. B-D: Overview of the sample acquisition and preparation for the data presented in this manuscript. The hearts of zebrafish larvae were temporarily arrested in a lightsheet chamber (B), and *z*-stack images acquired (C) which were pre-processed prior to use in *morphoHeart* (D). E-F: Snapshots of the *morphoHeart* GUI, showing the image segmentation tab (E), and the morphometric analysis tab (F).

Figure S2

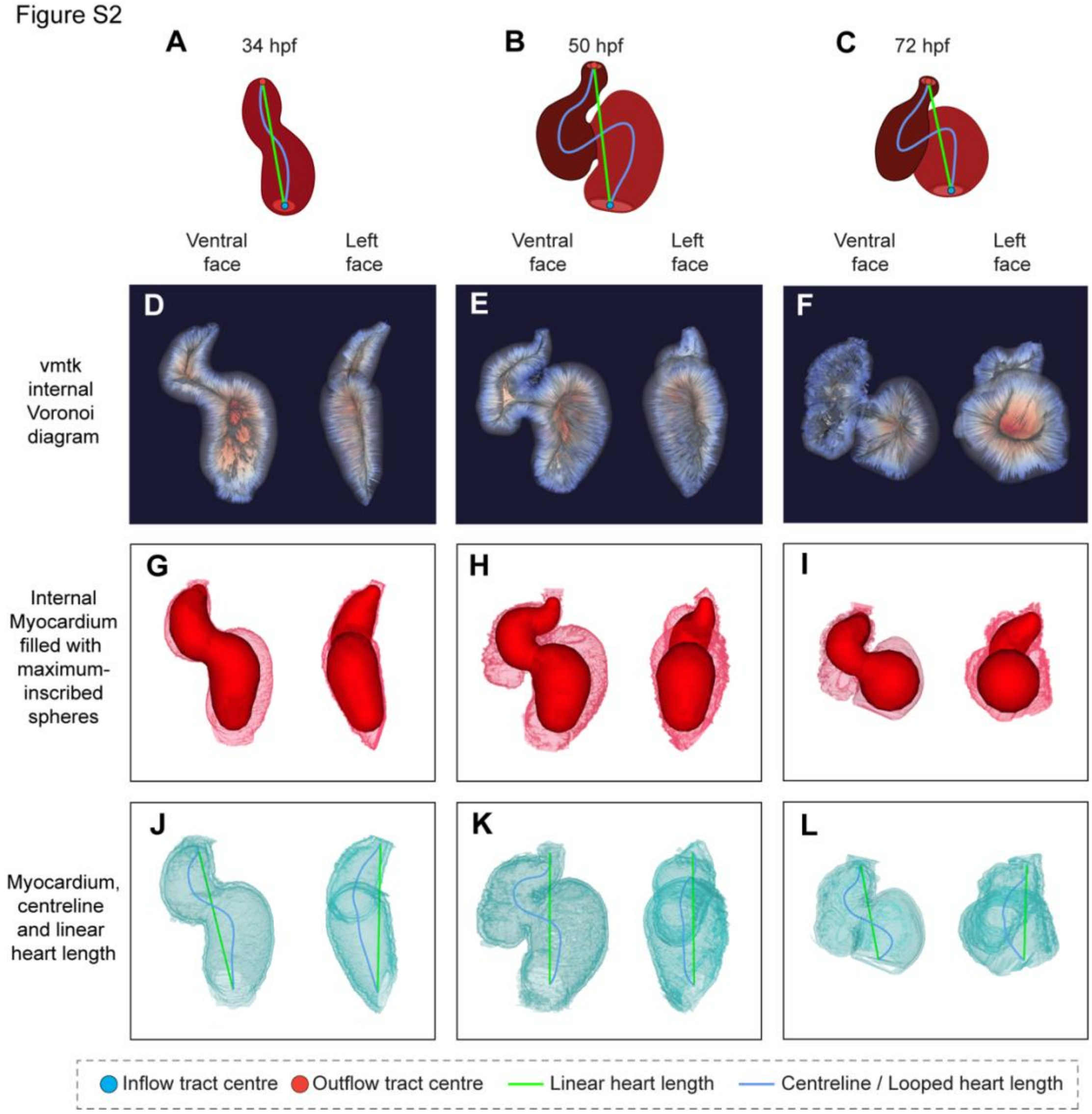


*Figure S2 - Acquisition of tissue 'centrelines'.*

A-C: Schematics depicting both linear (green) and looped (blue) lines through the heart between inflow and outflow poles. The blue looped line represents the 'centreline' through the tissue generated by assuming the heart as a tubular structure. D-E: Internal Voronoi diagram generated when extracting the centreline using the vmtk package. G-I: Internal myocardial mesh filled with maximum-inscribed spheres used to calculate each point of the centreline. J-L: Myocardial tissue meshes showing linear heart length (green, linear distance between poles) and looped heart length (blue, vmtk-calculated centreline through tissue between poles).

Figure S3

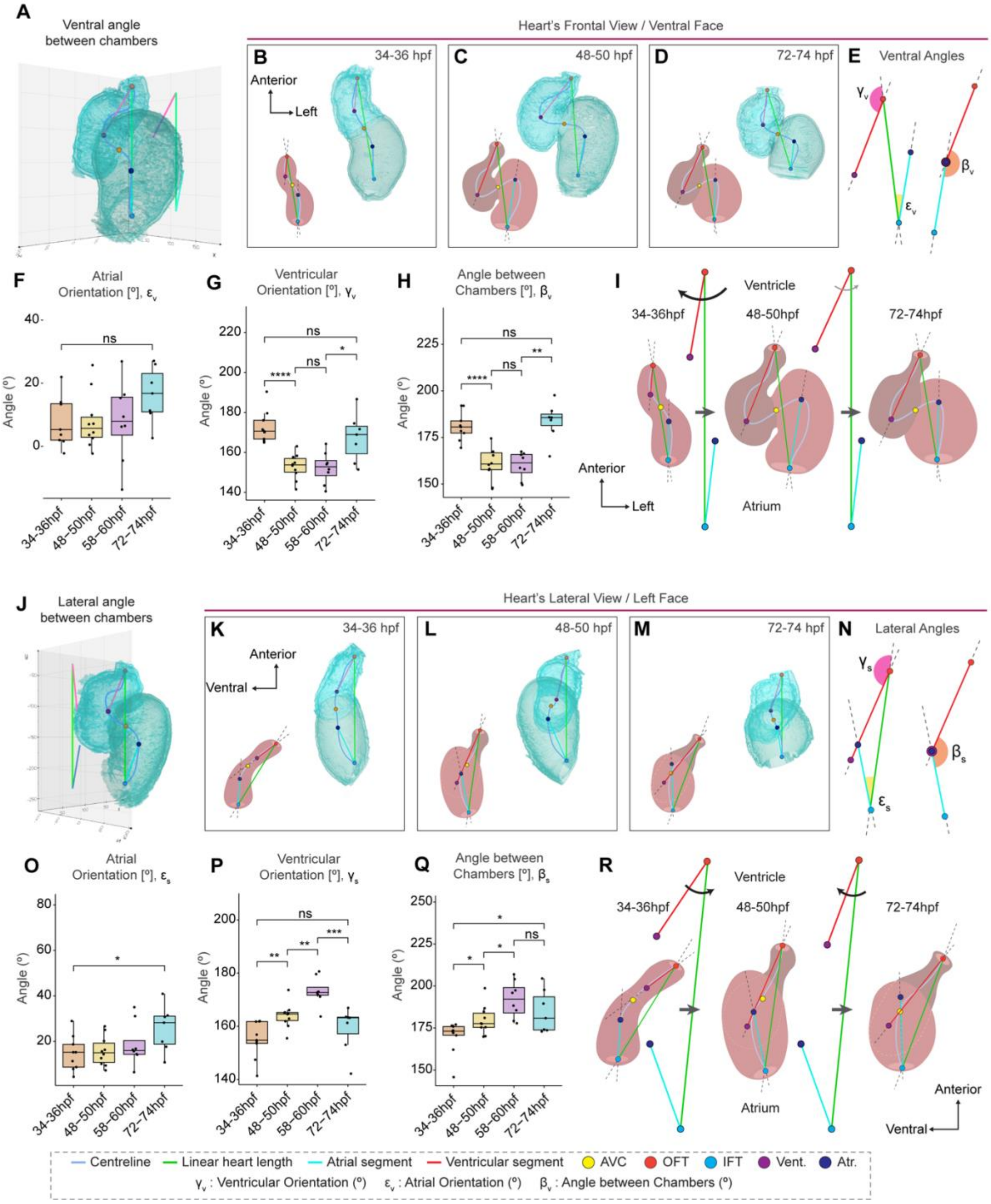


*Figure S3 - Heart morphogenesis is accompanied by relative chamber realignments.*

A-I: Schematic depicting method for measuring sagittal chamber orientations (A), and examples of chamber positions at each time point (B-D). Each chamber angle is measured, and the sagittal angle between them calculated (E). The atrium remains static (F), while the ventricle first moves substantially clockwise, and after looping pivots counter-clockwise (G), resulting in a sagittal displacement and realignment of the chambers as the heart loops, grows, and compacts (H, I). J-R: Schematic depicting method for measuring lateral chamber orientations (J), and examples of chamber positions at each time point (K-M). Each chamber angle is measured, and the lateral angle between them calculated (N). The atrium's lateral position remains relatively unchanged (O), while the ventricle straightens as the heart loops, and becomes laterally displaced as it compacts (R). This results in repositioning of the chambers or lateral rotation around the AVC (Q, R). One-way ANOVA with multiple comparisons.* $p<0.5$, ** $p<0.01$, *** $p<0.001$, **** $p<0.0001$. 34-36hpf: n=9; 48-50hpf: n=10; 58-60hpf: n=8; 72-74hpf: n=7.

Figure S4

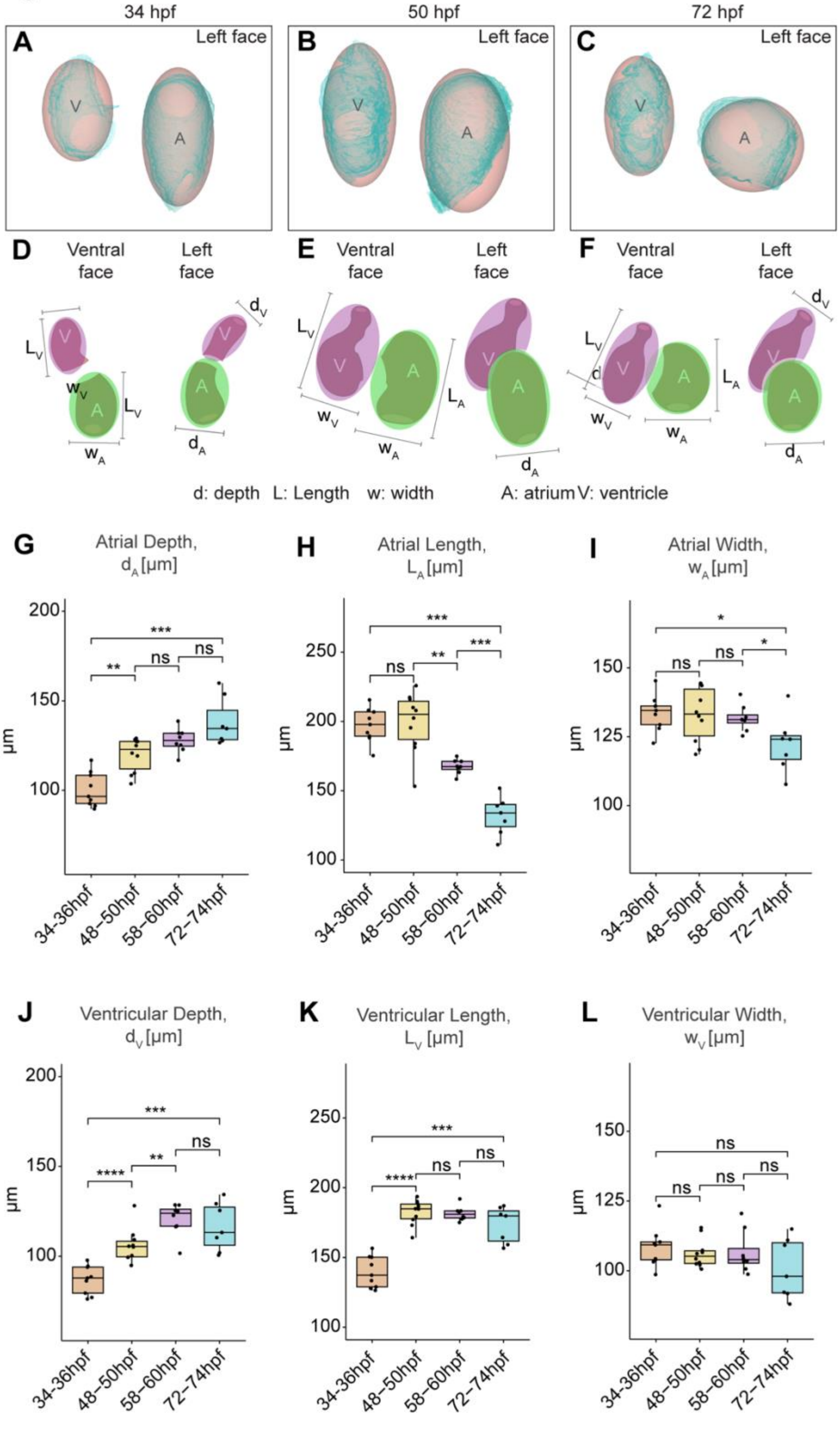


*Figure S4 - The chambers undergo different geometrical shape changes.*

A-C: Individual chamber meshes with fitted ellipsoids, lateral view from the left side. D-F: Illustration of the geometrical measurements acquired for both atrium (A) and ventricle (V), including chamber depth (d), length (L) and width (w), at each developmental stage. Both ventral view and lateral left-sided views are shown. G-L: Quantification of chamber depth (G,J), length (H,K) and width (I,L). The atrium expands in the z-plane, becoming deeper throughout development (G). Between 48hpf and 74hpf it also shortens (H) and narrows (I), becoming more spherical. The ventricle also expands in depth (J), whilst it lengthens (K) and narrows (L) between 34hpf and 50hpf to become a more elongated shape. One-way ANOVA with multiple comparisons.* $p<0.5$, ** $p<0.01$, *** $p<0.001$. 34-36hpf: n=9; 48-50hpf: n=10; 58-60hpf: n=8; 72-74hpf: n=7.

Figure S5

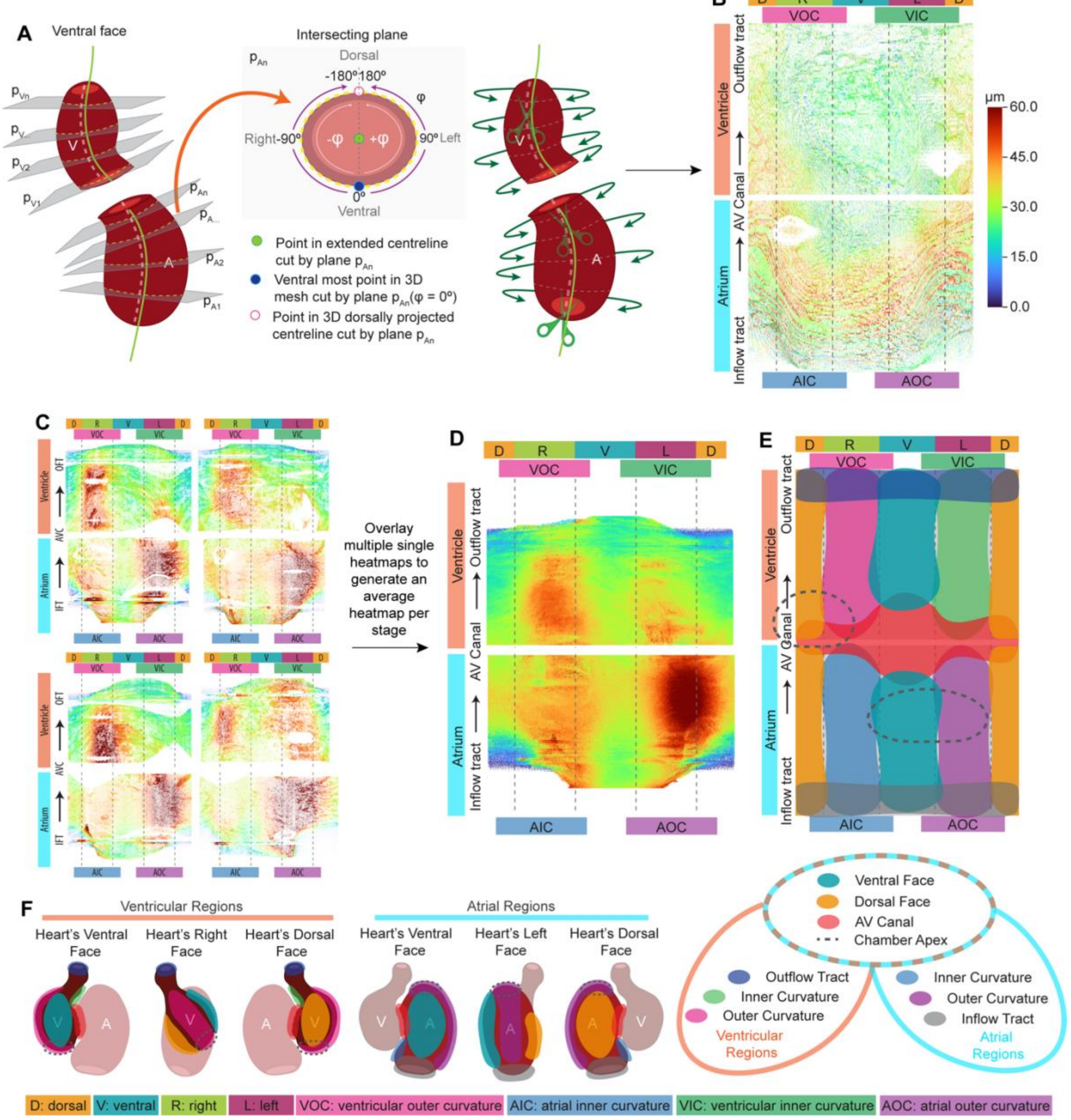


*Figure S5 - Generation of individual and averaged 2D heatmaps.*

A: Schematic depicting the method for unrolling 3D heatmaps into 2D heatmaps. The centreline (green) is extended beyond the poles of the heart, and a defined number of planes are cut through the heart mesh, transverse to the orientation of the centreline at each plane. For each intersecting plane, the ventral-most point of the mesh is cut and assigned position 0, while the dorsal-most point of the mesh is cut and assigned position 180/-180, giving each mesh point in each plane a universal coordinate, and which is associated with it's relevant thickness/ballooning value. This allows the tube to be 'unrolled', and mapped onto a standard 2D geometry (B). C-D: As each heatmap has the same coordinate system, the thickness/ballooning measurement can be averaged at each coordinate, allowing individual heatmaps from multiple embryos at the same stage and of the same genotype to be combined, producing an average heatmap which represents typical phenotype independent of small biological variation in tissue morphology (D). E-F: Schematics to facilitate interpretation of the heatmaps, demonstrating which regions of the 2D heatmaps relate to which morphological region in the heart. Labels around the heatmap indicate cardiac region: D - dorsal, V - ventral, L - left, R - right, AOC - atrial outer curvature, AIC - atrial inner curvature, VOC - ventricular outer curvature, VIC - ventricular inner curvature, AVC - atrioventricular canal.

Figure S6

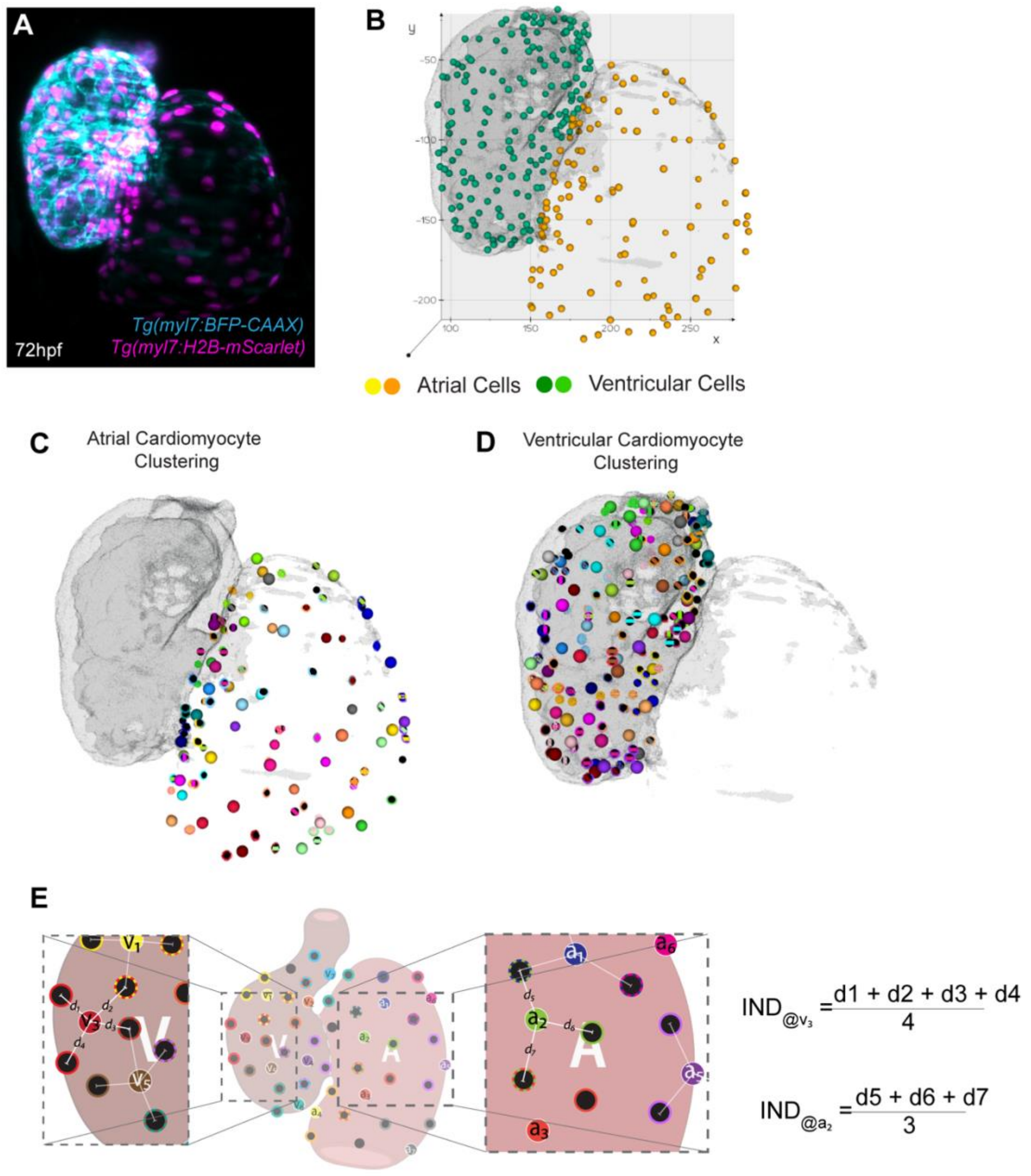


*Figure S6 - morphoCell module facilitates analysis of cell number and size.*

A: Maximum intensity projection of the heart of a *Tg(myl7:BFP-CAAX); Tg(myl7:H2B-mScarlet)* transgenic embryo, highlighting the myocardium (blue) and myocardial nuclei (magenta). Dots indicate nuclei identified using the Imaris spot-finder function. B-D: *morphoCell* renderings of myocardium and cardiomyocyte nuclei (spheres). Nuclei are initially not categorised into chambers (B), however chambers can be separated via a user-defined plane, and nuclei subsequently automatically categorised as atrial (orange spheres) or ventricular (green spheres). C-E: In each chamber, a subset of cells are automatically assigned as ‘seed’ cells (single colour) or ‘cluster’ cells (striped), for a defined number of clusters per seed. Seeds and clusters can be categorised to regions of interest within the chamber. The 3D distances between the seed cell and its cluster cells is measured, and averaged to generate a single average internuclear distance value for each seed.

Figure S7

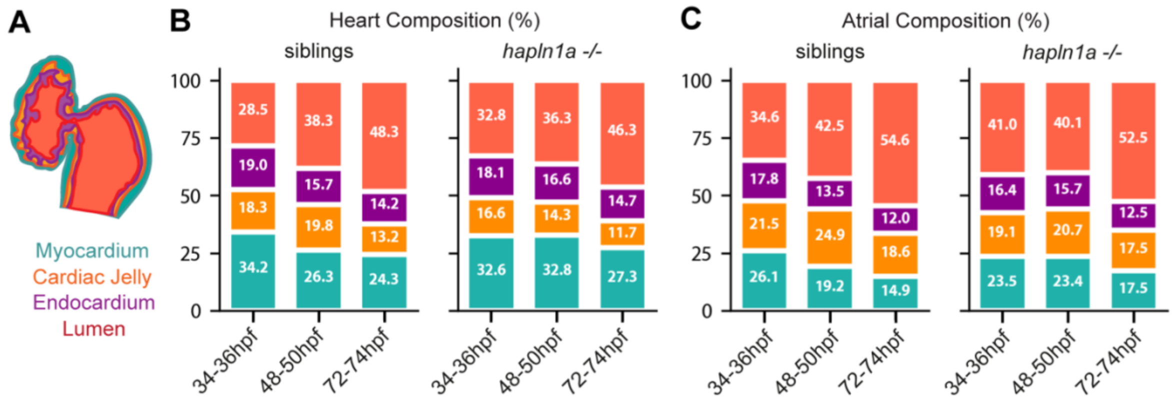


*Figure S7 - Relative ECM contribution to total heart volume is disrupted in hapln1a mutants.*

A-C: Analysis of heart composition by percentage contribution of each compartment of the heart. Schematic depicts contributing tissues/regions of the heart (A). During early stages of heart morphogenesis, *hapln1a* mutants display a comparative reduction in contribution of ECM to total heart or atrial volume (B,C).

**Pre-*morphoHeart* Masking and Cropping FIJI Macro**

```
/*
 *  merge_myoc_and_cj.ijm
 *  This macro will:
 *  - Ask the user to define a directory where the images are saved
 *  - Ask the user to open two individual channels of an image.
 *  - Create a MIP using both channels
 *  - Ask the user to create square in the MIP that includes all the ROI (heart) and use such
square
 *    to crop both channels
 *  - Ask the user to create a black square box to separate all signal from the borders (in
Ch1)
 *  - Use the black square to separate also the signal from Ch2.
 *  Then, in each channel the macro will:
 *  - Enhance contrast using 0.3% saturated pixels and normalizing the histogram
 *  - Enhance contrast equalizing the histogram
 *  - Despeckle (x3)
 *  - Save each cropped channel individually _EDC.tif
 *  Finally, for each channel the macro will create channel masks to use in morphoHeart
using the
 *  morphological filters from MorpholibJ
 *
 *  Created by:
 *  Juliana Sanchez-Posada
 *
*/

macro "ECD and mask" {
	print("\\Clear")
	print("NEW RUN");

	// Get images source directory
	dirSource = getDirectory("Choose the directory where the final channels will be
saved... ");
	print("dirSource: "+dirSource);

	run("ROI Manager...");

	//Open channel 1
	waitForUser("Open Channel 1 and click OK when ready.");
	ch1_ID = getImageID();
	ch1o_tt = getTitle();
	ch1_tt = split(ch1o_tt, '.')
	ch1_name = ch1_tt[0];
	ch1_filename = dirSource + ch1_name;
	print(ch1_ID, ch1o_tt, ch1_name);
	print(ch1_filename);
```

```
	//Open channel 2
	waitForUser("Open Channel 2 and click OK when ready.");
	ch2_ID = getImageID();
	ch2o_tt = getTitle();
	ch2_tt = split(ch2o_tt, '.');
	ch2_name = ch2_tt[0];
	ch2_filename = dirSource + ch2_name;
	print(ch2_ID, ch2o_tt, ch2_name);
	print(ch2_filename);

	//EDC for both channels
	// Ch1
	selectImage(ch1o_tt);
	fECandDx3();
	ch1_EDC = getImageID();
	ch1EDC_tt = getTitle();

	// Ch2
	selectImage(ch2o_tt);
	fECandDx3();
	ch2_EDC = getImageID();
	ch2EDC_tt = getTitle();

	//Merge channels
	Label_Comp = "c2=["+ch2EDC_tt+"] c5=["+ch1EDC_tt+"] create keep";
	run("Merge Channels...", Label_Comp);
	//Max Intensity Projection
	run("Z Project...", "projection=[Max Intensity]");
	MAX_ID = getImageID();
	MAX_tt = getTitle();
	saveAs("tif", dirSource+ch1_name+"_"+ch2_name+"_EDC Composite");

	// Set line width to 10
	run("Line Width...", "line=10");
	run("Colors...", "foreground=black background=black selection=yellow");

	//Crop image
	//Select MAX image to draw square to crop
	selectImage(MAX_ID);
	setTool("rectangle");
	waitForUser("Draw square -Shift- that encompasses the whole heart and click OK
when ready");
	roiManager("Add");

	//Crop MIP
	selectImage(MAX_ID);
	roiManager("Select", 0);
```

```
	run("Crop");
	saveAs("tif", dirSource+ch1_name+"_"+ch2_name+"_EDC Crop Composite");

	//Crop ch1
	selectImage(ch1_EDC);
	roiManager("Select", 0);
	run("Crop");
	setTool("rectangle");
	waitForUser("Draw rectangle that encloses the whole image and avoids open
contours and click OK when ready");
	if (selectionType() == 0) {
		roiManager("Add");
		run("Draw", "stack"); // same as Ctrl+D
		print("Black rectangle drawn in ch1");
	}
	makeRectangle(0, 0, 1, 1);
	saveAs("tif", ch1_filename+"_EDC");
	ch1ECD_ID = getImageID();
	ch1ECD_tt = getTitle();

	//Crop ch2
	selectImage(ch2_EDC);
	roiManager("Select", 0);
	run("Crop");
	roiManager("Select", 1);
	run("Draw", "stack"); // same as Ctrl+D
	print("Black rectangle drawn in ch2");
	makeRectangle(0, 0, 1, 1);
	saveAs("tif", ch2_filename+"_EDC");
	ch2ECD_ID = getImageID();
	ch2ECD_tt = getTitle();

	waitForUser("Check stacks and click OK when ready to continue");

	selectImage(ch1ECD_ID);
	masking();
	ch1mask_ID = getImageID();
	ch1mask_tt = getTitle();
	saveAs("tif", ch1_filename+"_mask");

	selectImage(ch2ECD_ID);
	masking();
	ch2mask_ID = getImageID();
	ch2mask_tt = getTitle();
	saveAs("tif", ch2_filename+"_mask");

	waitForUser("Check stacks and click OK when ready to close");
	close("*");
```

```
	roiManager("reset")

	close("*");
	close("\\Others");

	print("Images have been closed");
	print("DONE: merge_myoc_and_cj");

}

function fECandDx3() {
	run("Enhance Contrast...", "saturated=0.3 normalize process_all");
	run("Despeckle", "stack");
	run("Despeckle", "stack");
	run("Despeckle", "stack");
}

function masking() {
	run("Morphological Filters (3D)", "operation=Laplacian element=Cube x-radius=2 y-
radius=2 z-radius=2");
	run("Invert", "stack");
	run("Threshold...");
	setAutoThreshold("Li dark");
	waitForUser("Set threshold (DO NOT CLICK APPLY) and click OK when ready");
	run("Convert to Mask", "method=Li background=Dark black");
	run("Invert", "stack");
	run("Morphological Filters (3D)", "operation=Dilation element=Cube x-radius=1 y-
radius=1 z-radius=1");
}
```
